# Alterations of redox and iron metabolism accompany development of HIV latency

**DOI:** 10.1101/549014

**Authors:** IL Shytaj, B Lucic, M Forcato, JM Billingsley, S Bosinger, M Stanic, F Gregoretti, L Antonelli, G Oliva, CK Frese, A Trifunovic, B Galy, C Eibl, G Silvestri, S Bicciato, A Savarino, M Lusic

**Affiliations:** Heidelberg University Hospital and German Center for Infection Research, Heidelberg, Germany; University of Modena and Reggio Emilia, Modena, Italy; Emory University, Atlanta, GA, USA; Institute for High Performance Computing and Networking, ICAR-CNR, Naples, Italy; University of Cologne, Cologne, Germany; Division of Virus-Associated Carcinogenesis, German Cancer Research Centre, Heidelberg, Germany; Leibniz-Forschungsinstitut für Molekulare Pharmakologie, Berlin and Institute of Biology, Cellular Biophysics, Humboldt Universität zu Berlin, Berlin, Germany; Italian Institute of Health, Rome, Italy

**Author notes:** equal contribution (IL Shytaj, B Lucic).

**Keywords:** oxidative stress, HIV-1 latency, SIVmac, Nrf2, iron, transferrin receptor-1, promyelocytic leukemia protein, nuclear bodies

## Abstract

Metabolic alterations, such as oxidative stress, are hallmarks of HIV-1 infection. However, their influence on the development of viral latency, and thus on HIV-1 persistence during antiretroviral therapy (ART), have just begun to be explored. We analyzed omics profiles of *in-vitro* and *in-vivo* models of infection by HIV-1 and its simian homolog SIVmac. We found that cells survive retroviral replication by upregulating antioxidant pathways and intertwined iron import pathways. These changes are associated with remodeling of the redox sensitive promyelocytic leukemia protein nuclear bodies (PML NBs), an important constituent of nuclear architecture and a marker of HIV-1 latency. We found that PML is depleted in productively infected cells and restored by ART. Moreover, we identified intracellular iron as a key link between oxidative stress and PML depletion, thus supporting iron metabolism modulators as pharmacological tools to impair latency establishment.

## Introduction

Since the discovery of HIV persistence during antiretroviral therapy (ART), the activation status of T-cells, and thus their metabolic activity, have been recognized as a primary determinant of HIV latency (Williams & Greene 2007). Latency is the main obstacle to HIV-1 eradication and is characterized by persistence of integrated viral DNA that is transcriptionally silent, but replication competent. During therapy, integrated viral DNA is mainly found in memory CD4^+^ T-cells (Chomont et al. 2009; Hiener et al. 2017) where proviral reservoirs can become transcriptionally activated to trigger systemic infection upon ART discontinuation (Chun et al. 1998; Chun et al. 1999). Latency is established early upon infection *in vitro* (Chavez et al. 2015) and *in vivo* (Chun et al. 1998) but the transcriptional mechanisms and metabolic pathways regulating the transition between productive and latent infection remain poorly explored.

Several concepts have been proposed to explain the development of HIV latency, and, among them, the reversal of activated cells to a resting state after infection is broadly accepted (Siliciano & Greene 2011). Among the different factors involved in this virus/host interplay, oxidative stress and the cellular antioxidant response have been linked to productive HIV-1 infection (Gorrini et al. 2013) and to reactivation from latency (Shytaj et al. 2013). Oxidative stress is characterized by the generation and accumulation of byproducts of oxygen metabolism, reactive oxygen species (ROS), which can play a double faced toxic and/or regulatory role (Benhar et al. 2016). To survive and regulate the accumulation of ROS, cells employ a plethora of antioxidant mechanisms. In particular, the Nuclear Factor, Erythroid 2 Like 2 (Nrf2) master gene is a transcriptional factor that regulates the expression of multiple antioxidant genes, including key redox modulators such as glutathione (GSH), nicotinamide adenine dinucleotide phosphate (NAPDH), the thioredoxin/thioredoxin reductase (Trx/TrxR1) axis, the quinone detoxifying agent NAD(P)H dehydrogenase (quinone 1) (NQO1) (Furuya et al. 2016; Zhang et al. 2009) and several molecules involved in cellular iron metabolism (Gorrini et al. 2013). While therapeutic manipulation, based on either blocking antioxidant defenses to target latently infected cells or inhibiting oxidative stress to impair HIV-1 replication is currently under pre-clinical and clinical investigation (Benhar et al. 2016); NCT02961829), the study of the cellular antioxidant responses upon infection has led to conflicting results (Furuya et al. 2016; Zhang et al. 2009; Gill et al. 2014).

Intracellular iron metabolism could represent a possible link between viral replication and redox signaling. Inside cells, iron can be found in its ferric (Fe(III) or ferrous (Fe(II) forms, with the latter being reactive and capable of promoting oxidative stress through the Fenton reaction (Wang & Pantopoulos 2011). The storage and export of iron are directly controlled by Nrf2, which regulates the ferritin H (FTH-1) iron storage molecule as well as the ferroportin (SLC40A1) iron exporter (Kerins & Ooi 2018; Muckenthaler et al. 2017). Altered levels of the iron import protein, transferrin receptor 1 (TfR1, encoded by *TFRC*) have been reported upon HIV-1 infection, and increased or decreased iron levels have been associated with enhanced or reduced viral replication, respectively, albeit only in cell lines (Savarino, Pescarmona, et al. 1999; Savarino, Calosso, et al. 1999; Drakesmith & Prentice 2008). While the interplay between oxidative stress and iron metabolism has been well established, its role in the setting of HIV-1 infection and its impact on redox-sensitive cellular proteins regulating HIV-1 replication and latency has not been explored. In this regard, another player potentially joining HIV-induced oxidative imbalance and latency maintenance is the promyelocytic leukemia protein (PML), which is the main building block of nuclear bodies (NB). The biogenesis of PML NBs, which associate with the silent viral genome in resting CD4^+^ T cells (Lusic et al. 2013), depends on the oxidation process. Upon oxidative stress generation, NB shell formation is facilitated, and PML protein is subsequently sumoylated and subjected to degradation (reviewed in (Sahin, de Thé, et al. 2014)). While we identified the proximity of silent proviral genomes to the PML NBs as a factor that contributes to maintenance of HIV-1 latency (Lusic et al. 2013), the question of how the virus enters the latent phase and whether PML NBs play a role during this process remains open.

While several redox-effects on HIV expression/latency are known, a general picture on the sequence of redox-related events accompanying HIV latency are still lacking, and data have often been derived from tumor cell line models for latent HIV infection. Here we use omics data from a primary CD4^+^ T-cell model of HIV latency and *ex-vivo* transcriptomic profiles derived from rhesus macaques infected with the HIV homolog SIVmac (Palesch et al. 2018) to identify a pathway unraveling through HIV-1 replication, oxidative stress, iron import and PML NBs depletion/reformation. This pathway characterizes the transition between the productive and latent phase of HIV-1 infection and highlights potential new druggable targets for therapeutic applications.

## Results

### Antioxidant response is upregulated during productive *in vitro* and *in vivo* infection

To study the expression of antioxidant genes and proteins during the different stages of HIV-1 infection, we used a primary CD4^+^ T-cell model similar to those previously adopted by several groups (Bosque & Planelles 2009; Lusic et al. 2013; Martins et al. 2016). In this model, CD4^+^ T-cells are activated through stimulation of CD3/CD28 receptors, infected with HIV-1_NL4-3_, and monitored over time until returning to a resting state (**Figure 1A**). This time course mimics distinct features of HIV-1 infection *in vivo, i.e.* a rapid initial growth of viral replication (**Figure 1B** and **1C**) accompanied by cell death or establishment of a small pool of cells harboring integrated viral DNA that can be reactivated (**Figure 1D, 1E** and **1F**). Donor-matched mock infected controls are used to standardize gene and protein expression levels, while sampling of mRNA and proteins at different time points enables specific analysis of the various stages of infection. To elucidate the relationship between viral replication and oxidative stress, we conducted RNA-Seq (**Figure 2**) and proteomic analyses (**Figure S1**) of a macro-pathway including a large body of genes (n=401) related to oxidative stress. We observed enriched expression of antioxidant genes and proteins in infected compared to mock infected control cells. Interestingly, the peak of enrichment was observed at days 7-9 post-infection (henceforth, dpi) (**Figure S1** and **2A**), corresponding to the time interval during which HIV-1 replication reaches its zenith (**Figure 1B** and **1C**). Consistently, the antioxidant response decreases alongside establishment of latency. Indeed, when we focus our analyses on infected cells and normalize the data to the latent time point (14 dpi), we observe enriched expression of antioxidant genes during productive infection (**Figure 2B**).

**Figure 1.**
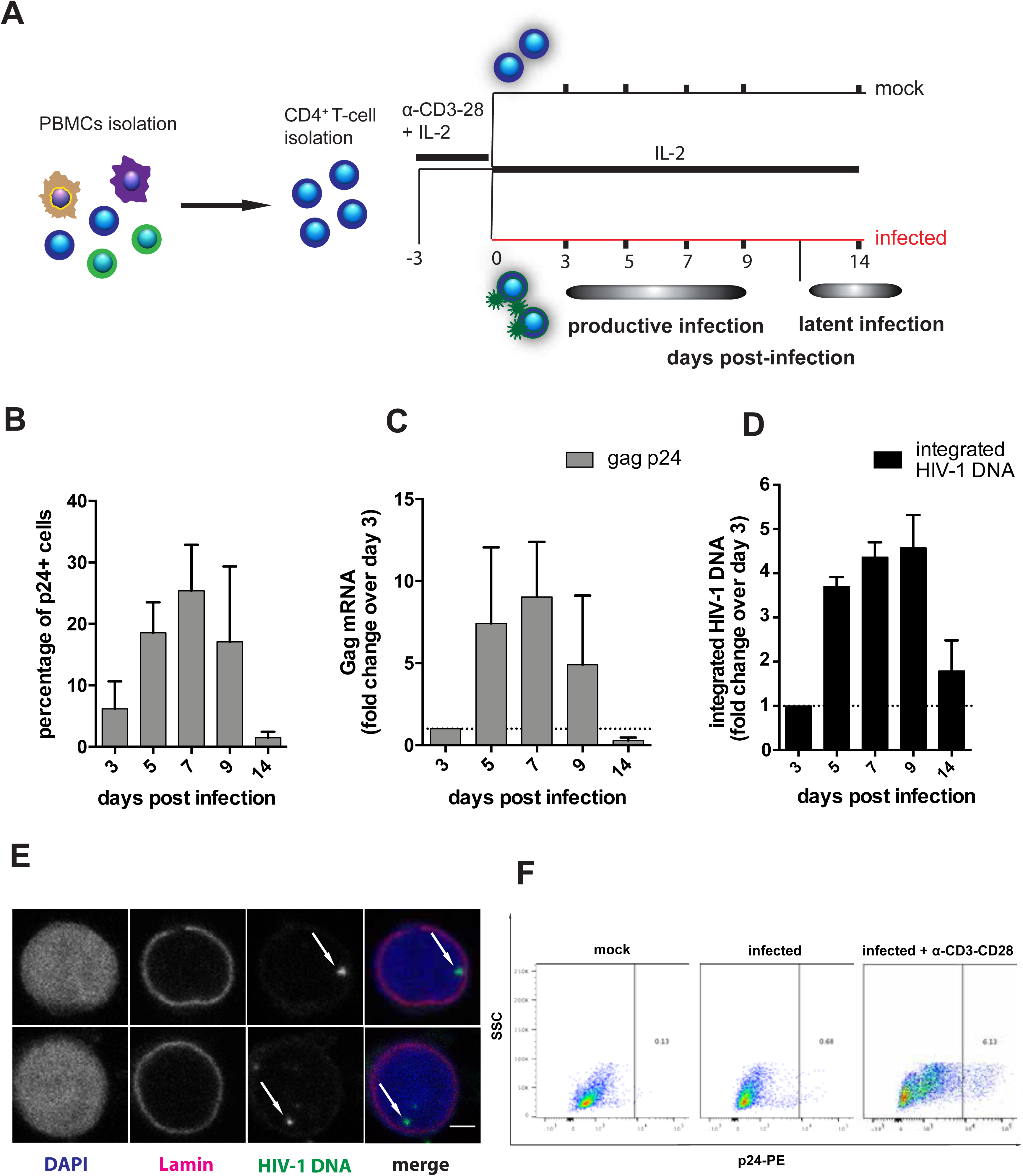
Primary CD4^+^ T-cell model of productive and latent HIV-1 infection. (A) Experimental design. (B) Percentage of intracellular gag p24^+^ CD4^+^ T-cells as measured by flow cytometry. (C) Relative levels of HIV-1 gag mRNA copies as measured by qPCR. (D) Relative levels of integrated HIV-1 DNA copies as measured by Alu-HIV PCR. (E) FISH detection of HIV-1 DNA at latency, *i.e.* 14 dpi (green: HIV-1; red: lamina). Scale bar = 2μm. (F) Production of gag p24 after 48 hr reactivation with α-CD3/CD28 beads at 14 dpi. Data in panels A-C are expressed as mean±SEM of 3 donors.

**Figure 2.**
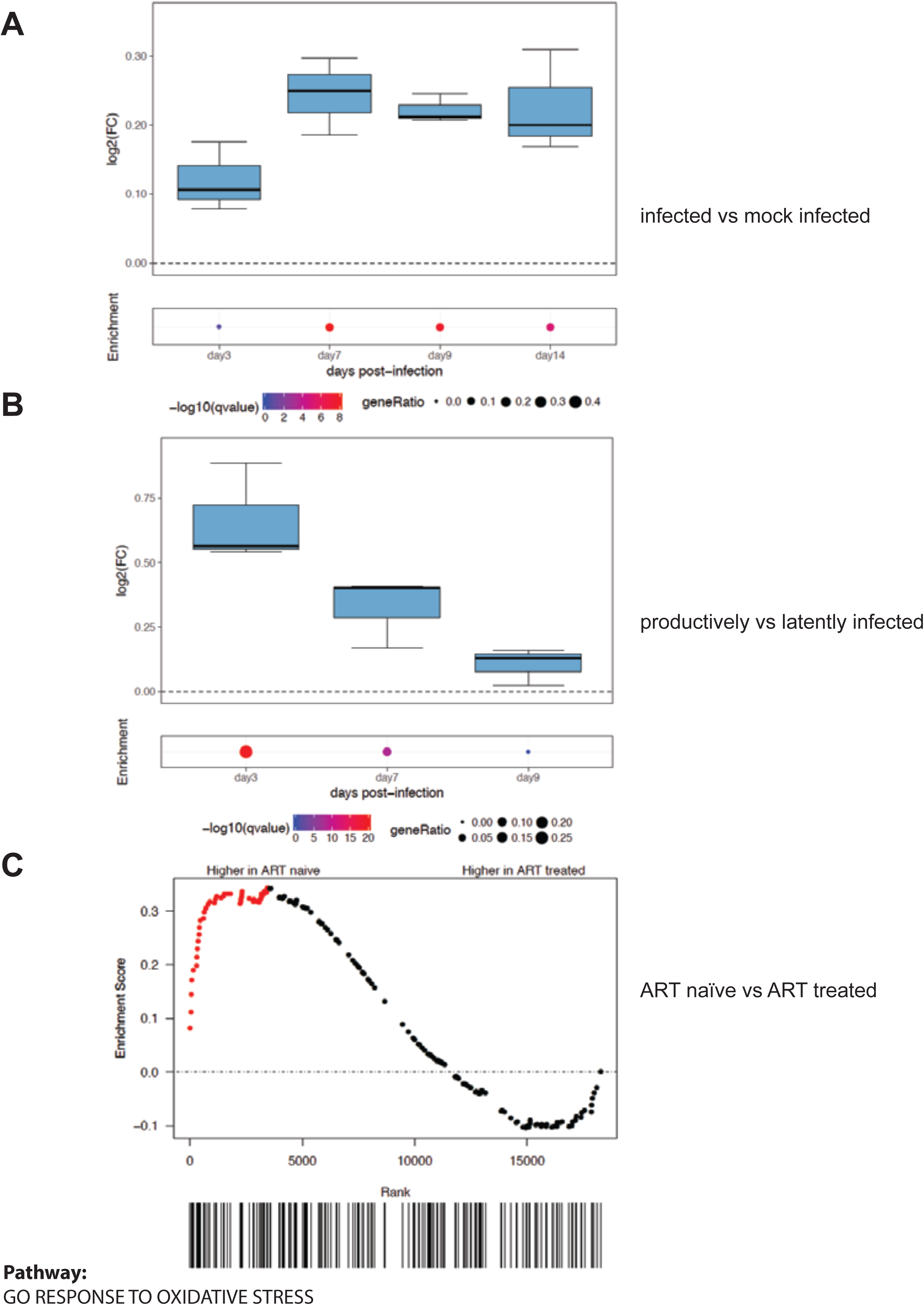
Viral replication induces upregulation of antioxidant genes *in vitro* and *in vivo*. RNA-Seq analysis of: different time points of primary CD4^+^ T-cells infected *in vitro* with HIV-1 or mock infected (panels A,B) and PBMCs of macaques infected with SIVmac239 before and after suppression of viremia with ART (panel C). Expression levels are normalized as: expression in infected vs matched mock-infected controls (panel A); expression in productively infected cells (3-7-9 dpi) vs expression in latently infected cells (14 dpi, panel B); expression in ART-treated vs ART naïve (panel C). For panels (A,B) number of donors =3, for panel C number of animals =8. For all panels the analyses were conducted on the the genes of pathway *GO response to oxidative stress* (401 genes; GO:0006979).

To investigate the *in-vivo* relevance of our findings we further analyzed the expression of genes involved in oxidative stress response using a recently generated RNA-Seq dataset from an animal model closely recapitulating the main features of HIV infection (Evans & Silvestri 2013; Palesch et al. 2018). This dataset is derived from peripheral blood mononuclear cells (PBMCs) of macaques chronically infected with the HIV homolog virus SIVmac in the presence or absence of ART. The use of this animal model provides an optimal *in-vivo* parallel of our *in-vitro* system, in that macaques can be standardized for viral inoculum, time/route of infection, and time points of analysis, with each animal acting as its own internal control before ART initiation (Evans & Silvestri 2013). In agreement with our *in-vitro* data, a Gene Set Enrichment Analysis (GSEA) showed enriched expression of antioxidant genes before the administration of ART (**Figure 2C**).

In conclusion, both *in-vitro* and *ex-vivo* data show that transcriptional and protein expression of antioxidant genes is enriched during the productive stage of viral infection and that this effect is reversed, at least in part, upon transition to a resting/latent state.

### HIV-1 replication drives nuclear translocation of Nrf2 and activation of its downstream antioxidant pathways

The transcriptomic and proteomic profiling of infected CD4^+^ T-cells points to a progressive enrichment of antioxidant defenses peaking with HIV-1 replication (**Figure S1** and **Figure 2**). This is further supported by a significant decrease in the reduced to oxidized glutathione (GSH/GSSG) ratio in infected versus mock infected control cells (mock: 53.0±8.5, infected: 38.9±9.2; mean±SEM; *P*= 0.021, paired *t*-test, data not shown), which is indicative of oxidative stress (Giustarini et al. 2013) and is consistent with the previously reported generation of oxidative species upon infection (Pace & Leaf 1995).

To characterize the antioxidant response upon HIV-1 infection we focused on the master transcription factor Nrf2 and its downstream antioxidant pathways. We first analyzed the subcellular localization of Nrf2, which is expected to relocalize from the cytosol to the nucleus upon activation (Ma 2013). We found that productive HIV-1 infection is accompanied by Nrf2 nuclear translocation, as shown by biochemical fractionation (**Figure 3A**). Moreover, single molecule HIV-1 RNA FISH combined with immunofluorescence (IF) for Nrf2 showed that the nuclear content of Nrf2 was higher in cells actively producing the virus (**Figure 3B**). We next analyzed the expression of six representative Nrf2-target genes involved in the utilization, synthesis or detoxification of redox species such as GSH, NAPDH, thioredoxin (Trx), iron and quinones (Gorrini et al. 2013). We observed a similar pattern of expression across all biological replicates (**Figure S2A**), with upregulation of all Nrf2 target genes during productive infection, both at the transcriptional (**Figure 3D**) and translational (**Figure 3E**) level. In all cases, a trend towards normalization to mock infected control levels was observed upon establishment of latency (**Figure 3D** and **3E)**. Moreover, a two-way ANOVA test highlighted the different time points (and thus stages) post-infection as the main source of variability (*P* <0.0001), irrespective of the gene analyzed (**Figure 3D**). These results suggest a broad upregulation of the Nrf2-regulated pathways driven by the increase in HIV-1 replication.

**Figure 3.**
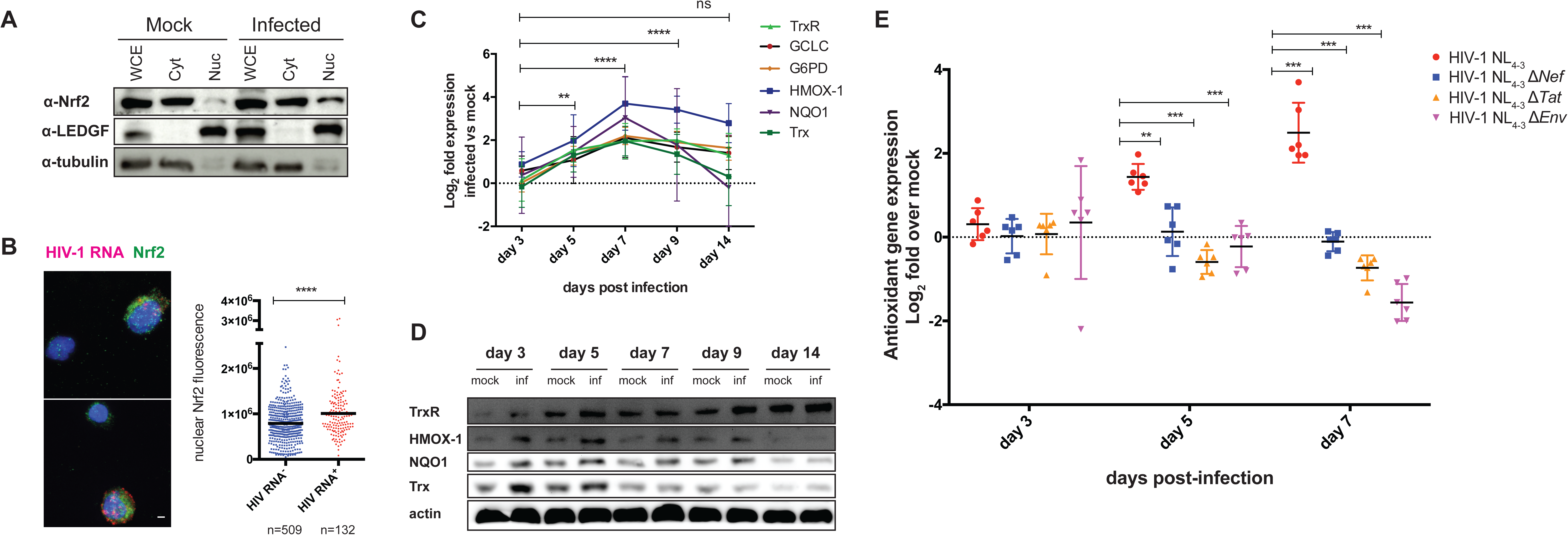
Activation of Nrf2 drives antioxidant responses to HIV-1 replication in CD4^+^ T-cells. (A) Subcellular localization of Nrf2 in CD4^+^ T-cells infected with HIV-1 or mock infected (3 dpi) as analyzed by biochemical fractionation. (B) Nuclear localization (left) and content (right) of Nrf2 in HIV-1 RNA^+^ and HIV-1 RNA^-^ cells (7 dpi) as measured by combining IF and HIV-1 RNA FISH. Scale bar = 2μm. n= number of cells from 3 donors. (C, D) Time course of the relative (infected vs mock infected) mRNA (C) and protein (D) levels of main targets of Nrf2 during the transition from productive (3-9 dpi) to latent (14 dpi) infection as measured by qPCR and western blot, respectively. TrxR1 and HMOX-1 were probed upon membrane stripping. (E) Comparison of the effect of different HIV-1 mutations on the mean relative (infected vs mock infected) mRNA level of the Nrf2 targets described in panel D. Values shown in panel B were calculated as nuclear corrected total cell fluorescence (as in(McCloy et al. 2014)) and analyzed by two-tailed unpaired *t*-test. For panels C, E, raw data were first normalized using GAPDH as housekeeping control and then expressed as Log_2_ fold mRNA expression in infected vs mock infected cells (calculated using the 2-ΔΔCTmethod (Livak & Schmittgen 2001). In panel E the average of the six genes listed in Panel C is shown. For both panels data were analyzed by-two way ANOVA followed by Turkey’s post-test for multiple comparisons. ** *P*<0.01; \*\*\**P*<0.001; \*\*\*\**P*<0.001. Trx= thioredoxin; NQO1= NAD(P)H Quinone Dehydrogenase 1; HMOX-1= Heme Oxygenase 1; G6PD= glucose-6-phosphate dehydrogenase; GCLC= Glutamate—cysteine ligase; TrxR1= thioredoxin reductase 1.

Previous studies have linked the production of ROS to specific HIV-1 proteins, such as gp120 (Pietraforte et al. 1994) and reverse transcriptase (Isaguliants et al. 2013), and to accessory proteins, in particular Nef (Vilhardt et al. 2002) and Tat (Gu et al. 2001; Zhang et al. 2009).

To assess how single viral proteins modulate the redox status of the cell in the context of infection, we employed three different viral clones bearing mutations in the *env (Pizzato et al. 2007), nef (Trautz et al. 2016)* or *tat (Bejarano et al. 2019)* genes. Compared to *wild type* HIV-1_NL4-3_, all mutant viruses exhibit the expected decrease in viral transcription (**Figure S2B**) and integration (**Figure S2C**), with the Δ*tat* mutant displaying the lowest fitness (**Figure S2B** and **S2C**). Of note, none of the mutant viruses triggers the upregulation of antioxidant genes (**Figure 3F**) when compared to their respective mock infected controls. Furthermore, as viral replication increases (5-7 dpi), the expression of antioxidant genes in cells infected with the *wild type* virus is always higher than in cells infected with either of the mutant viruses (**Figure 3F**).

Overall, these data demonstrate that HIV-1 replication induces oxidative stress, followed by activation of Nrf2 and upregulation of its downstream pathways. Moreover, our results suggest that efficient viral replication, rather than single viral proteins, is required to activate the Nrf2 pathway.

### Upregulation of Nrf2 targets and transition to latency are associated with increased iron import

Iron metabolism is a key target of Nrf2 signaling (Gorrini et al. 2013). Our data highlighted heme oxygenase-1 (HMOX-1), the enzyme responsible for the catabolism of heme leading to ferrous iron (Fe(II)) release (Gozzelino et al. 2010), as one of the antioxidant factors highly upregulated in the early infection stages (**Figure 3D** and **3E**). To have a broader view of iron metabolism regulation during the different stages of infection, we analyzed the expression of several iron homeostasis genes in our omics data sets. RNA-Seq and proteomics data derived from our *in-vitro* model revealed enriched expression of mRNA and proteins regulating iron metabolism (**Figure 4A**, left panel and **Figure S3A** left panel). The *ex-vivo* data derived from SIV-infected macaques further showed two sets of iron metabolism genes differentially enriched either before or during ART (**Figure 4A**, right panel). As iron homeostasis can be regulated by mutually counteracting mechanisms (such as import and export) we narrowed our analyses to iron import regulation (Muckenthaler et al. 2017). Both our *in-vitro and ex-vivo* data indicate enriched expression of iron import genes towards the transition to latency or after ART administration (**Figure 4B**). In addition, the two iron transport proteins that could be detected in our proteomic analysis showed a similar trend (**Figure S3A**, right). TfR1 was the iron import marker that could be readily detected in all our omics data sets. We thus verified its expression independently using flow cytometry (FACS) and IF (**Figure 4C** and **4D** and **Figure S3B**). We observed an early decrease in TfR1 expression 3 dpi, most likely secondary to intracellular iron release through the action of HMOX-1 (**Figure 4C**). As viral replication progresses (7-9 dpi), an opposite pattern is observed (**Figure 4C** and **4D** and **Figure S3B**), showing enhanced iron import capacity via TfR1 internalization (**Figure 4D**) in infected cells during viral production and upon transition to latency. Absolute expression of TfR1 is decreased upon transition to a resting state (14dpi, **Figure 4C** and **Figure S3B**), in line with the low expression of this marker in resting lymphocytes (**Figure S3C**). Yet, infected cells still display higher TfR1 expression compared to mock infected (**Figure 4C**), supporting the omics data and suggesting a persistent imbalance of iron metabolism in infected cells upon latency establishment. Consistently with the expression pattern of TfR1, FTH-1 and SLC40A1 expression was increased early upon infection suggesting enhanced iron storage and export, an effect that was reversed in the later stages (**Figure 4E**). Further supporting the notion that increased iron import sustains viral replication, the addition of non toxic levels (**Figure S3D**) of the iron donor FeCl_3_ 6H_2_O to the culture medium of infected cells was found to enhance gag p24 production.

**Figure 4.**
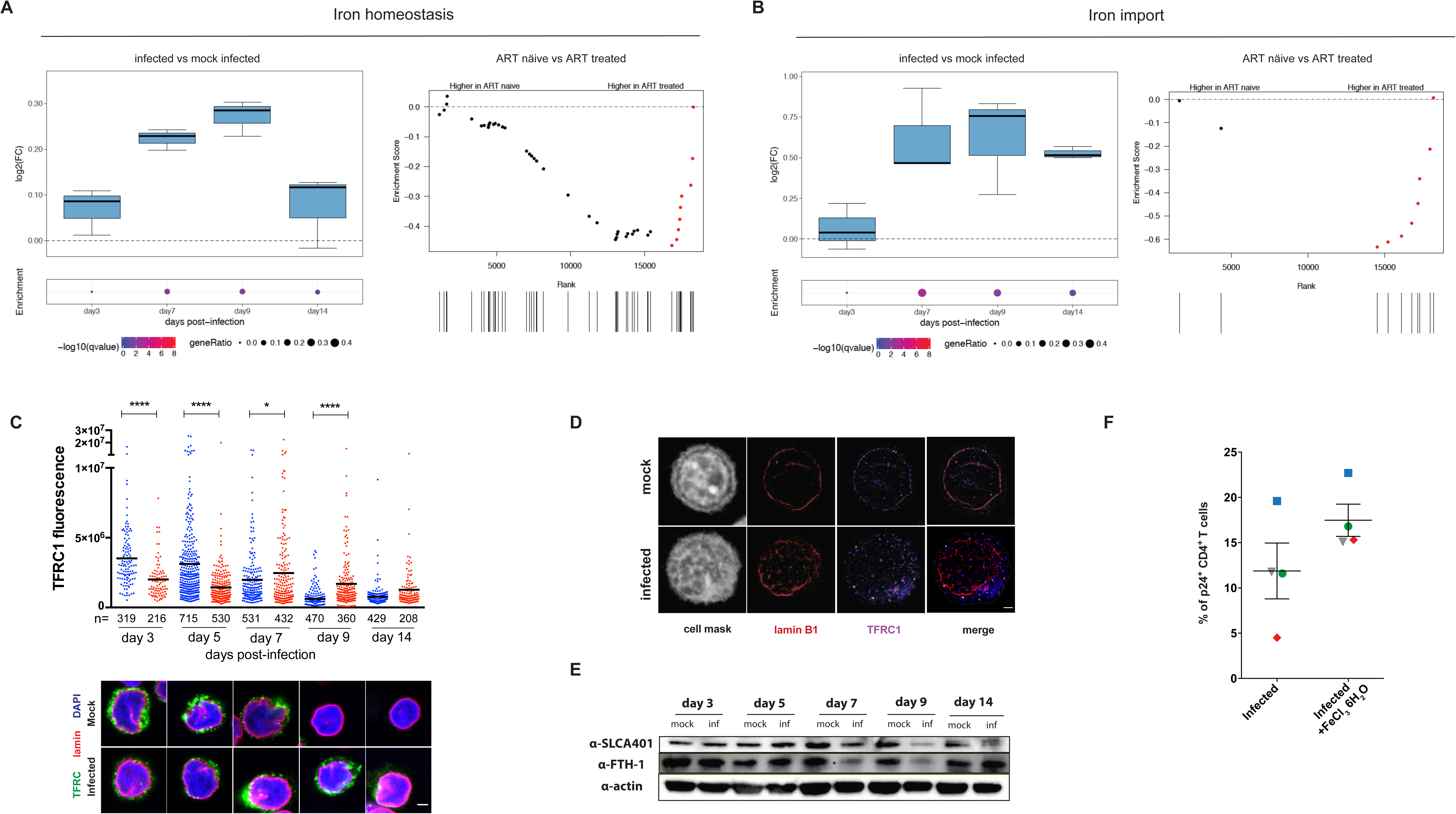
HIV-1 replication enhances iron consumption and import during transition to latency. (A,B) RNA-Seq analysis of: different time points of primary CD4^+^ T-cells infected *in vitro* with HIV-1 or mock infected (left panels) and PBMCs of macaques infected with SIVmac239 before and after suppression of viremia with ART (right panels). The gene sets used for the analyses are *GO cellular ion iron homeostasis* (46 genes, GO:0006879; panel A) and *GO iron ion import* (12 genes, GO:0097286; panel B). (C,D) TfR1 expression (C) and subcellular localization (D) in HIV-1 infected or mock infected CD4^+^ T-cells as visualized by confocal and STED microscopy, respectively. Fluorescent values shown in panel C were calculated as described in Figure 3 and in (McCloy et al. 2014). Data were analyzed by one-way ANOVA followed by Sidak’s post-test to compare corresponding mock infected vs infected pairs (mean±SEM; n= number of cells from 3 donors). (E) Protein expression over time of FTH-1 and SLC40A1 in HIV-1 infected or mock infected CD4^+^ T-cells as assessed by western blot. (F) Percentage of intracellular p24^+^ gag CD4^+^ T-cells infected with and left untreated or treated for 48 hr with the iron donor FeCl_3_ 6H_2_O at 150μM concentration. Scale bars = 2μm. \**P*<0.05; \*\*\*\**P*<0.001.

Overall, these data show that intracellular iron levels are distinctively altered during the different stages of HIV-1 infection, with progressively increased iron import during the transition to latency.

### HIV-1 replication induces degradation of the redox-sensitive latency marker PML

We then tested the impact of the above described metabolic imbalances on the host cells and the course of the infection. For this we examined the PML protein, which is redox-sensitive (Sahin, de Thé, et al. 2014; Niwa-Kawakita et al. 2017) and colocalizes with latent HIV-1 DNA (Lusic et al. 2013). First, we analyzed the regulation of the PML pathway (including PML and its main interacting proteins) during the different stages of HIV-1 infection (**Figure 5A**) by comparing HIV-1 infected to mock infected cells. Transcriptional enrichment of the PML pathway was observed in infected cells, peaking during productive infection (**Figure 5A**); a similar trend was observed in *ex-vivo* samples from SIVmac infected macaques (albeit without reaching statistical significance **Figure 5B**). Conversely, protein levels displayed the highest enrichment in infected cells only at the latent time point (**Figure 5C**).

**Figure 5.**
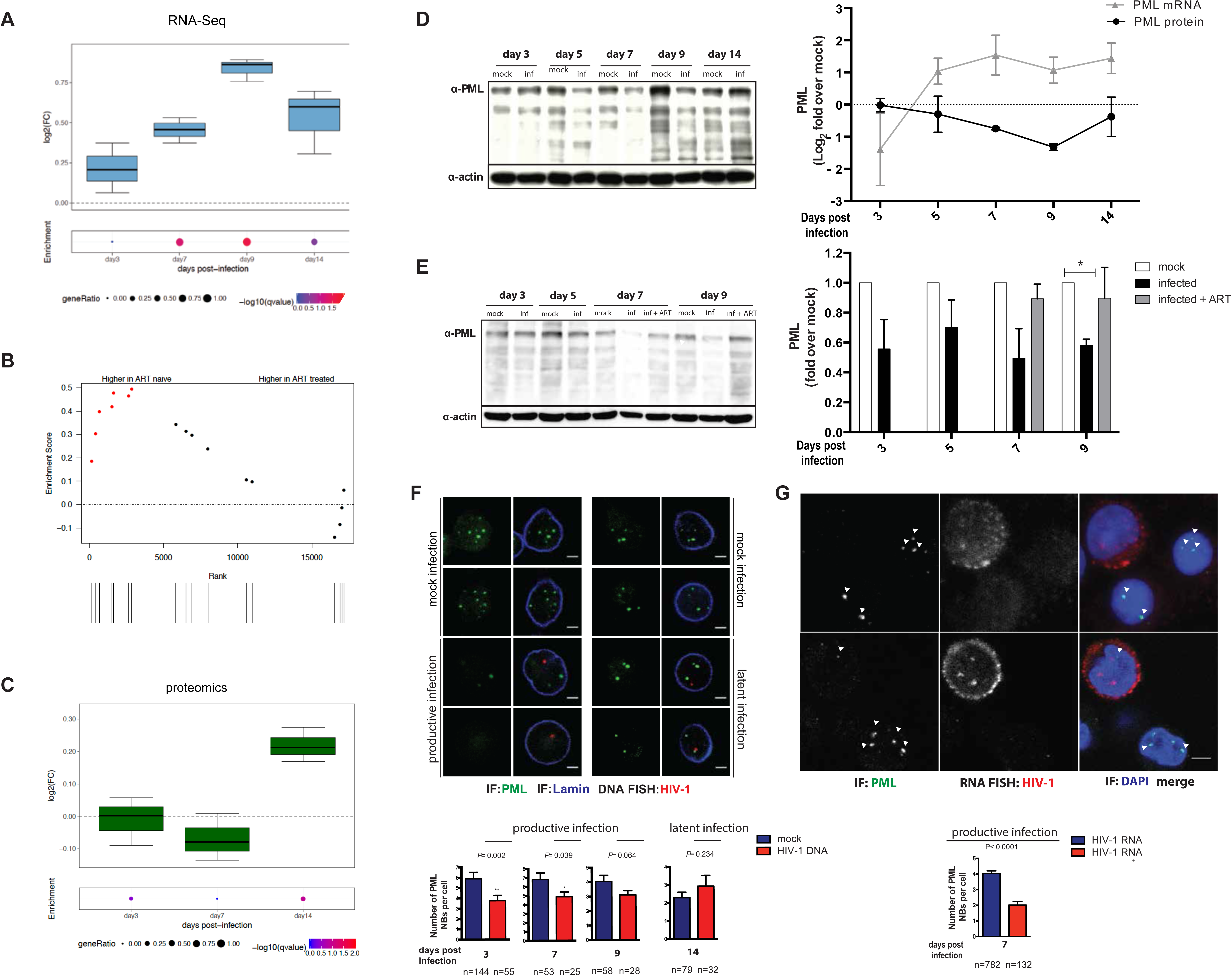
Interplay between the redox-sensitive marker PML and HIV-1 expression. (A,B) RNA-Seq analyses of different time points in primary CD4^+^ T-cells infected *in vitro* with HIV-1 or mock infected (A) or PBMCs of macaques infected with SIVmac239 before and after suppression of viremia with ART (B). (C) Proteomic analysis of HIV-1 infected and mock infected CD4^+^ T-cells. The gene set considered for all analysis is *Biocarta_PML_pathway* (17 genes, M4891) (D) Western blot (left panel) and relative expression (right panel) of PML protein and mRNA in HIV-1 infected vs mock infected CD4^+^ T-cells over time. (E) Western blot (left panel) and relative expression (right panel) of PML protein over time in mock infected vs HIV-1 infected CD4^+^ T-cells with or without ART (added at 5 dpi). (F,G) Representative image and quantification of the number of PML bodies in HIV-1 DNA^+^ vs mock infected cells (panel F) and HIV-1 RNA^+^ vs HIV-1 RNA^-^ (panel G) CD4^+^ T-cells. n= number of cells from 2 (panel F) and 3 (panel G) donors. Scale bar = 2μm. For panel D and E western blot data were quantified using Fiji-Image J (Schindelin et al. 2012) and all data were normalized over an housekeeping protein (beta-actin, western blot) or gene (GAPDH, qPCR). For microscopy analyses infected cells were identified with either DNA (panel F) or RNA (panel G) FISH and PML NBs were stained by IF. The algorithm used for automatic PML counting is detailed in the materials and methods section. Data were analyzed by two-tailed unpaired *t*-test.

To explore the specific modulation of PML only, we analyzed its expression by qPCR and western blotting (**Figure 5D,** left and right panels, respectively). We observed a progressive decrease in the amount of PML protein alongside the increase in viral replication (5-9 dpi), followed by a partial replenishment of PML upon transition to latency (14 dpi) (**Figure 5D,** left panel). Of note, the level of PML mRNA displayed an opposite trend, suggesting a compensatory increase to counteract ongoing protein degradation (**Figure 5D**, right panel). The trend displayed by PML was opposite to that observed for antioxidant defenses (**Figure 3C** and **3D**). In line with this, uninfected CD4^+^ T-cells treated with the potent pro-oxidant H_2_O_2_ showed rapid nuclear import of Nrf2 (**Figure S4A**) and decreased levels of PML (**Figure S4B**), albeit associated with high cellular mortality.

To estimate the impact of HIV-1 replication on PML depletion, we suppressed viral production using ART (5 dpi). The addition of ART abrogated p24 production in less than 48 hours (data not shown) and restored PML protein to the mock infected control levels (**Figure 5E**).

To exclude potential biases due to bystander effects, which are typical of oxidative stress (Klammer et al. 2015), we performed single cell analysis of cultures harboring HIV-1 nucleic acids. By combining HIV-1 DNA FISH with IF for PML NBs (**Figures 5F**), we confirmed degradation of PML during productive HIV-1 infection as well as its reformation in cells harboring latent HIV-1 (**Figure 5F**). Furthermore, single cell HIV-1 RNA FISH and IF staining for PML NBs proved that CD4^+^ T-cells productively infected with HIV-1 are characterized by lower numbers of PML NBs (**Figure 5G**). Identifying 3D objects, such as PML bodies, by projecting z-stacks in 2D can potentially bias the counting of such objects. Hence, we used an algorithm (described in materials and methods) that automatically generates 3D cell reconstructions and identifies HIV-1 RNA and PML NBs signals. The results confirmed significantly lower numbers of PML bodies in cells actively transcribing HIV-1 RNA (**Figure S4C** and **S4D**). Finally, as PML NBs have been previously described as possible restriction factors for HIV-1 replication (Dutrieux et al. 2015; Turelli et al. 2001; Kahle et al. 2015) we depleted PML with arsenic trioxide (As_2_O_3_), (**Figure S5A, S5B** and **S5C**) and infected the cells with either *wild type* HIV-1_NL4-3_ or with the orange and green HIV-1 (OGH) dual color reporter virus that allows the discrimination between productive and latent infection based on expression of fluorescent markers (Vranckx et al. 2016; Calvanese et al. 2013). The data showed increased viral production upon PML depletion (**Figure S5D** and **S5E**). Viral integration/latency, on the other hand, displayed variable trends (**Figure S5D** and **S5E**), suggesting that PML depletion through As_2_O_3_ favors productive over latent HIV-1 infection (**Figure S5D**).

Overall these results show that PML is degraded by HIV-1 replication and suggest that the interplay between HIV-1 and PML might be both a cause and a consequence of viral production.

### Iron mediates PML depletion upon HIV-1 infection

Both As_2_O_3_ and iron (in particular in its ferrous state) can lead to increased ROS production, potentially influencing the stability of PML. We thus tested the effect of a non toxic (Ciciliano et al. 2015) (**Figure S3D**) concentration of the iron donor FeCl_3_ 6H_2_O on PML NBs. We found that iron per se could induce PML and PML NBs depletion, recapitulating the effects of HIV-1 infection (**Figure 5D**). Of note, the effect or iron is lower compared to As_2_O_3_ (**Figure 6A, 6B** and **6C**). Treatment with As_2_O_3_ has been previously associated with altered subcellular distribution of PML in cell lines, including increased cytoplasmic localization (Hands et al. 2014). We used STimulated Emission Depletion (STED) microscopy to investigate the subcellular localization of PML NBs in primary CD4^+^ T-cells 48 hr after exposure to As_2_O_3_ or FeCl_3_ 6H_2_O, respectively. However, we could not detect increased cytoplasmic PML NBs localization in any of the conditions examined (**Figure 6D**).

**Figure 6.**
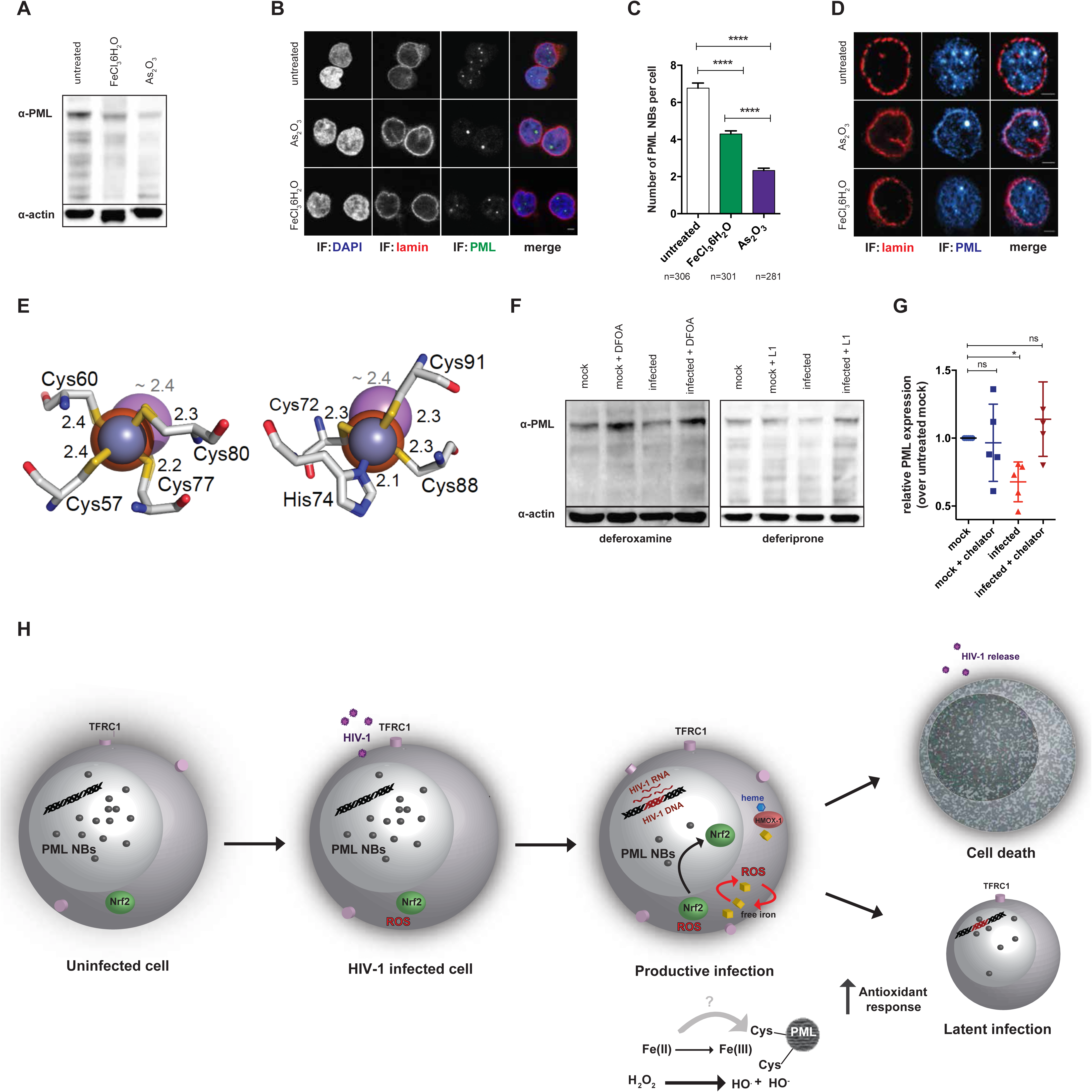
Influence of iron on PML stability. (A-C) PML and PML NBs expression in CD4^+^ T-cells left untreated or treated for 48 hr with 5μM As_2_O_3_ or 500μM FeCl_3_ 6H_2_O as measured by western blot (A) or IF (B,C). n= number of cells from 3 donors. Scale bar 2μm. (D) Subcellular localization of TfR1 in HIV-1 or mock infected CD4^+^ T-cells at 7 dpi as visualized by STED microscopy. Scale bar 2μm. (E) Structural models of the coordination, in the PML-ring domain (pdb code: 5yuf), of the canonical binding of Zn2+ (blue sphere) compared to Fe(II) (orange sphere) and As3-(purple sphere). Coordinating residues of zinc finger sites 1 (left panel) and 2 (right panel) are shown. Binding distances are indicated in Å (black for Fe(II) and Zn(II) and grey for As(III)). The predicted binding of Fe(III) overlaps with that of Fe(II) and it is excluded for simplicity. (F,G) PML expression in CD4^+^ T-cells infected with HIV-1 or mock infected at 5dpi as measured by western blot. Cells were left untreated or treated with the iron chelator deferoxamine (DFOA, 1μM) or deferiprone (L1, 50μM). (H) Schematic depiction of the regulation of antioxidant responses, intracellular iron content and PML NBs abundance during the transition between productive and latent HIV-1 infection. The PML NBs numbers depicted in C were calculated using automatic algorithms as detailed in the materials and methods section. Western blot quantifications in (G) were done with Fiji-Image J (Schindelin et al. 2012), normalized over the housekeeping protein beta-actin and expressed as fold change over untreated, mock infected cells. Data, expressed as mean±SEM, were analyzed by two-tailed unpaired *t*-test (C; 3 donors) or one-way ANOVA followed by Dunnet’s multiple comparisons test (G; 5 donors).

In addition to its action as pro-oxidant, it has been previously demonstrated that PML degradation by As_2_O_3_ is at least in part mediated by arsenic replacing zinc through direct binding to the cysteine residues of the RING fingers located in the RING finger, B-box, and coiled-coil (RBBC) domain of PML (Zhang et al. 2010). Similarities between the atomic radius of iron and zinc, and between the oxidation state of Fe(III) and As_2_O_3_, suggest a possible direct effect of iron on PML through replacement of zinc. We thus used a recently published high resolution crystal structure of PML RING domain tetramer to replace the zinc ion bound to PML with iron (Wang et al. 2018). Our molecular modelling suggests that iron could in principle replace zinc in the RING finger (**Figure 6E**). However, the model showed that, unlike As_2_O_3_, the binding of either Fe(II) or Fe(III) does not alter the coordination of the cysteine residues nor the Fe-S binding length. Hence, these results suggest that iron alters PML stability mainly through its pro-oxidant effect.

To specifically investigate the role of iron as a mediator of PML depletion induced by HIV-1, we tested the effects of two iron chelators: deferoxamine (DFOA) and deferiprone (L1). Both compounds have been shown to possess antiviral activity (Georgiou et al. 2000). To exclude potential bias and isolate the effect of iron chelation we selected non-toxic (Figure S6A) concentrations of DFOA and L1 that could reduce intracellular iron levels (as assessed by TfR1 upregulation, **Figure S6B**), but did not impair HIV-1 replication (**Figure S6C**). Because long-term incubation with iron chelators decreased the viability of infected cells (**Figure S6D**) (Georgiou et al. 2002), we analysed cells 5dpi, corresponding to a time point where PML is already depleted while cell viability is preserved. We found that incubation with either DFOA or L1 could revert PML depletion in infected cells, while leaving mock infected cells unaffected (**Figure 6F** and **6G**).

Overall, these data prove that iron is sufficient and necessary to induce PML depletion in the context of HIV-1 infection, and suggest that the effect of iron on PML is exerted indirectly through enhancement of oxidative stress.

## Discussion

Our study shows that dysregulation of cellular redox and iron metabolism is specifically driven by HIV-1 replication in CD4^+^ T-cells. Increasing oxidative stress and enhancing iron intake, leading to ROS generation, HIV-1 reshapes important components of the nuclear architecture, PML NBs, by causing depletion of PML protein and ultimately favoring further viral production. This self-sustaining mechanism is counteracted by a potent antioxidant response of the host, peaking concomitantly to maximal viral replication and preluding to latency establishment (**Figure 6H**).

We show that primary CD4^+^ T-cells surviving at peak of viral replication display enhanced iron-import capacity through TfR1. TfR1 upregulation in infected cells is likely associated with the utilization of iron for the activity of ribonucleotide reductase (Drakesmith & Prentice 2008), which is essential for sustaining both cellular and viral replication. It is tempting to speculate that upregulation of TfR1 selects cells that can survive the increased iron consumption imposed by the infection, and therefore harbour latency. This hypothesis is in line with a previous study conducted in tumor cell line models, showing that downregulation of TfR1 is associated with death of HIV-infected cells (Savarino, Pescarmona, et al. 1999). This is likely due to the fact that TfR1 downregulation, which in healthy cells limits iron overload, leads, in infected cells, to levels of iron intake that are unable to sustain both HIV-1 and cellular replication. In line with this model, the iron chelator L1 induced selective killing of the infected cell cultures. The increased iron requirement of infected cells is also supported by the persistence of increased iron import capacity upon development of latency *in vitro* and *in vivo* in macaques.

Iron binds TfR1 in its ferric (Fe(III) form, but, once imported in cells, is converted in endosomes to its reactive ferrous form Fe(II) (MacKenzie et al. 2008). In the oxidized environment caused by HIV-1 infection, Fe(II) can in turn enhance ROS production through the Fenton reaction (**Figure 6H**). Evidence for an enhanced oxidative stress in our model is derived from the consumption of reduced glutathione (GSH), an intracellular antioxidant tripeptide which acts as a ROS scavenger (reviewed in (Benhar et al. 2016)).

Noteworthy, the role of iron in regulating the stability of the redox-sensor PML protein is here proven. Molecular modelling suggests that iron could replace zinc in the RING finger of PML, but that its PML-degrading effect is, unlike that of As_2_O_3_, probably indirect, and exerted through increased oxidative stress. Accordingly, we show that the potent ROS H_2_O_2_, in line with previous studies (Sahin, Ferhi, et al. 2014), is able to deplete PML in CD4^+^ T-cells. The timing of maximal PML depletion during infection (7-9 dpi) further corroborates an indirect effect of iron, in that it overlaps with the upregulation of TfR1 in infected cells and consequently with the increased presence of the redox active Fe(II). Even though the binding of iron to the PML RING finger domain is not predicted to destabilize its structure with an arsenic-like mechanism, the spatial proximity resulting from this binding could increase the susceptibility to oxidation of the cysteines in the domain, as previously described for As_2_O_3 (Zhou et al. 2015)_. Future studies will be required to test this hypothesis.

It was previously shown that PML depletion increases stability and nuclear translocation of the antioxidant master transcription factor Nrf2 (Guo et al. 2014; Niwa-Kawakita et al. 2017). The antioxidant response that we observed in the present study is case in point, in that it was associated with broad activation of Nrf2 downstream targets. Moreover, the pattern of PML depletion mirrored that of the expression of Nrf2 targets, suggesting that degradation of PML might contribute to the generation of the potent antioxidant response induced by HIV infection.

Several studies have examined the independent role of single HIV-1 proteins in influencing the redox status of the cell (Pietraforte et al. 1994; Isaguliants et al. 2013; Vilhardt et al. 2002; Zhang et al. 2009; Gu et al. 2001). While we could not ascribe the generation of antioxidant responses to any specific viral protein, our data show that these responses develop only in the presence of efficient and sustained replication of *wild type* HIV-1. In line with this, all the mutants tested in this study displayed antioxidant responses comparable to mock-infected cells. Previous studies have described both activation (Zhang et al. 2009) and inhibition of the Nrf2-regulated pathways upon HIV-1 infection (Gill et al. 2014). Cell type differences might explain these discrepancies, in that inhibition of Nrf2 has been reported in myeloid cells, in particular in the central nervous system (Gill et al. 2014). Antioxidant responses were decreased upon latency establishment in our model, but this did not lead to their full normalization, as compared to mock-infected cells. In line with this, a previous study indicated upregulation of antioxidant defenses in latently infected cell lines as compared to their uninfected counterparts (Bhaskar et al. 2015). Similarly, enhanced antioxidant responses were associated with inhibition of pathways inducing HIV-1 RNA transcription, such as those mediated by NF-kappaB and NF-AT (Bosque & Planelles 2009; Benhar et al. 2016). On the whole, this evidence suggests that viral replication might exert a selective pressure based on the antioxidant capacity of infected cells. According to this model, the strength of the antioxidant response would determine the ability of the cell to survive the infection while favoring the development of latency. Alternatively, low-level ongoing viral replication might cause the persistence of residual oxidative stress, and of the cellular response. Intriguingly, the permanent upregulation of antioxidant defenses in latently HIV-1 infected cells, parallels a condition observed in cancer cells (Benhar et al. 2016). Pre-clinical and clinical studies suggest that this persistent upregulation might render HIV-infected and cancer cells more susceptible to pharmacologically induced oxidative bursts, namely through the inhibition of the Nrf2-targets TrxR1 and GCLC (Benhar et al. 2016).

Finally, reformation of PML upon latency supports the possible role of PML NBs as a marker of silent HIV-1 DNA, as previously shown (Lusic et al. 2013). We could distinctively highlight both degradation and reformation of PML in cells harboring HIV-1 DNA, thus proving the specificity of the effect aside of the inevitable bystander effects that characterize redox metabolism. Reformation of PML NBs was concomitant to the transition to a resting state, suggesting that indirect mechanisms, such as mitigation of TfR1 upregulation in resting cells (MacKenzie et al. 2008), might facilitate the reduction of oxidative stress and consequently the normalization of PML levels in infected cells.

By combining omics data sets and single cell analyses, the present study contributes to a general picture of the interplay between redox and iron metabolism in the setting of HIV-1 infection. It presents, though, some limitations such as circumscribing the analyses to bulk CD4^+^ T-cells or PBMCs to allow for longer post-infection follow-up. Different T-cells subsets are, however, characterized by variable baseline antioxidant defenses (Chirullo et al. 2013) and could thus contribute to a different degree to the effects that we observed. Of note, a recent study has elegantly characterized the metabolic activity of T-cell subsets and shown the correspondence with their different predisposition to HIV-1 infection (Valle-Casuso et al. 2018). Another potential limitation of our study is the combined analysis of all different isoforms of PML, which can independently influence its function and localization (Condemine et al. 2006). Further studies will be required to investigate if specific isoforms of PML might be differentially influenced by HIV-1 infection.

Overall, the present work proves that HIV-1 can alter the nuclear structure of the host and favor its own replication by increasing oxidative stress and intracellular iron import, while the cellular response to these effects preludes to latency development. Moreover, these data open the possibility of combining drugs to increase oxidative stress and iron content to induce PML degradation and target the HIV-1 reservoir.

## Materials and Methods

### Primary CD4^+^ T-cell isolation and culture

Whole blood of healthy donors was provided by Heidelberg University Hospital Blood Bank following approval by the local ethics committee. CD4^+^ T-cells were isolated using the RosetteSep™ Human CD4^+^ T Cell Enrichment Cocktail (STEMCELL Technologies Inc., Vancouver, British Columbia, Canada) following manufacturer’s instruction. After isolation cells were activated by adding the Dynabeads® Human T-Activator CD3/CD28 using a bead/cell ratio of 1:2. Cells were cultured in RPMI 1640 supplemented with 20% fetal bovine serum (FBS), penicillin/streptomycin, and 10 ng/mL IL-2 and kept at a concentration between 1-2 *10^6^ cells/mL in an incubator at 37°C in a 5% CO_2_ atmosphere. Three days after activation cells were infected and/or treated with drugs as described below.

### *In-vitro* infection with HIV-1

For *in-vitro* infection, 5-20*10^6^ activated CD4^+^ T-cells were used. Cells were pelleted and infected using either *wt* HIV-1_NL4-3_ (2ng p24/10^6^ cells) the *nef* mutated virus described in (Trautz et al. 2016) (10ng p24/10^6^ cells; HIV-1_NL4-3_ Δ*nef*), the dual color OGH HIV-1 virus (Calvanese et al. 2013) (1μg p24/10^6^ cells), the *tat* mutated virus described in (Bejarano et al. 2019) (1μg p24/10^6^ cells; HIV-1_NL4-3_ Δ*tat*) or *env* mutated virus pseudotyped with vesicular stomatitis virus glycoprotein (Pizzato et al. 2007) (1μg p24/10^6^ cells; HIV-1_NL4-3_ Δ*env*). For infection with the *wt* HIV-1_NL4-3_ or HIV-1_NL4-3_ Δ*nef* virus, cells were left 2-4 hr in an incubator at 37°C and 5% CO_2_. For infection with the OGH virus, HIV-1_NL4-3_ Δ*tat* and HIV-1_NL4-3_ Δ*env*, cells were spinoculated at 2300rpm for 1.5 hr at 37°C. Following infection cells were washed and resuspended at 1*10^6^ cells/mL in RPMI 1640 supplemented with 20% FBS and 10 ng/mL IL-2.

### Drug treatments

For testing pharmacologically-induced PML depletion or iron overload/chelation CD4^+^ T-cells at day 3 post-activation were left untreated, or treated with 1) arsenic trioxide (As_2_O_3_; Sigma Aldrich, Saint Louis, MI, USA, ref: 356050) at 5μM concentration, 2) iron(III) chloride hexahydrate (FeCl_3_ 6H_2_O; Sigma Aldrich, Saint Louis, MI, USA, CAS: 10025-77-1) at 500μM, 3) Deferoxamine mesylate salt (DFOA, Sigma Aldrich, Saint Louis, MI, USA, CAS: 138-14-7) at 1μM concentration 4) Deferiprone (L1, Sigma Aldrich, Saint Louis, MI, USA, CAS: 30652-11-0) at 50μM concentration. Cells treated with FeCl3 6H_2_O were harvested 48 hr post-treatment and used for western blot analyses. Cells treated with As_2_O_3_ and DFOM were infected and used for further analyses as described below. For specifically blocking HIV-1 replication, CD4^+^ T-cells were incubated with a three drug ART five days post-infection with *wt* HIV-1. The antiretroviral combination was composed of T-20 (10μM), raltegravir (10nM) and efavirenz (100nM) (kindly provided by the National Institutes of Health AIDS Research and Reference Reagent Program).

### RNA-Seq analyses

RNA sequencing was performed on an Illumina Next Seq platform and CASAVA was used to perform base calling. The quality of the 2×75 bp reads was assessed with FastQC (https://www.bioinformatics.babraham.ac.uk/projects/fastqc/). STAR (Dobin et al. 2013) version 2.5 was used to align raw reads to build version hg38 of the human genome and to calculate counts for GENCODE (release 25) basic annotated genes. Normalization and differential analysis were carried out using the DESeq2 R package (Love et al. 2014). Genes were considered differentially expressed with an adjusted p-value lower than 0.05.

Gene sets representing the activity of pathways were derived from the Molecular signature database (Liberzon et al. 2011). The statistical over-representation of gene sets among differentially expressed genes and proteins was assessed with a one-tailed Fisher exact test. R (version 3.3.1) was used for statistical analyses.

### Proteomic analysis

Sample preparation was performed using the Single-Pot Solid-Phase-enhanced Sample Preparation approach SP3, as described elsewhere (Hughes et al. 2019). In brief, 2 µL of a 1:1 mixture of hydrophilic and hydrophobic carboxylate coated paramagnetic beads (SeraMag Speed Beads, #44152105050250 and #24152105050250, GE Healthcare, Little Chalfont, UK) were added to 30 µg protein of each sample. Acetonitrile was added to achieve a final concentration of 50% organic solvent. Bound proteins were washed with 70% ethanol and 100% acetonitrile. Beads were resuspended in 5 µL 50 mM Triethylammoniumbicarbonate buffer containing 0.6 µg Trypsin (SERVA) and 0.6 µg LysC (Wako). Digestion was carried out for 16 hr at 37°C in a PCR cycler. Recovered peptides were resuspended in 1% formic acid / 5% DMSO and stored at 20°C prior MS analysis. All samples were analyzed on a Q-Exactive Plus (Thermo Scientific) mass spectrometer that was coupled to an EASY nLC 1200 UPLC (Thermo Scientific). Peptides were loaded with solvent A (0.1% formic acid in water) onto an in-house packed analytical column (50 cm × 75 µm I.D., filled with 2.7 µm Poroshell EC120 C18, Agilent) equilibrated in solvent A. Peptides were chromatographically separated at a constant flow rate of 250 nL/min using the following gradient: 3-5% solvent B (0.1% formic acid in 80 % acetonitrile) within 1 min, 5-30% solvent B within 91 min, 30-50% solvent B within 17 min, followed by washing at 95% for 10 min. For library generation the mass spectrometer was operated in data-dependent acquisition mode. The MS1 survey scan was acquired from 350-1300 m/z at a resolution of 70,000. The top 10 most abundant peptides were isolated within a 2 Th window and subjected to HCD fragmentation at a normalized collision energy of 27%. The AGC target was set to 5e5 charges, allowing a maximum injection time of 55 ms. Product ions were detected in the Orbitrap at a resolution of 17,500. Precursors were dynamically excluded for 20 s. A data-independent acquisition method was employed for protein quantification. The mass spectrometer was operated in data-independent acquisition (DIA) mode. The MS1 scan was acquired from 400-1220 m/z at a resolution of 140,000. MSMS scans were acquired for 10 DIA windows at a resolution of 35,000. The AGC target was set to 3e6 charges. The default charge state for the MS2 was set to 4. Stepped normalized collision energy was set to A, B, C = 23.5%, 26%, 28.5%. The MSMS spectra were acquired in profile mode. Data analysis: The raw data of the pooled library samples were processed with Maxquant (version 1.5.3.8) using default parameters. Briefly, MS2 spectra were searched against the Uniprot human database, including a list of common contaminants. False discovery rates on protein and PSM level were estimated by the target-decoy approach to 1% (Protein FDR) and 1% (PSM FDR) respectively. The minimal peptide length was set to 7 amino acids and carbamidomethylation at cysteine residues was considered as a fixed modification. Oxidation (M) and Acetyl (Protein N-term) were included as variable modifications. The match-between runs option was disabled. For DIA quantification raw data were processed with Spectronaut Pulsar X (version 11) using default parameters. Briefly, MS2 spectra were searched against the previously generated library. The maximum of major group Top N was set to 6, the decoy method to inverse and the data filtering was set to Qvalue.

### Measurement of reduced (GSH) and oxidized (GSSG) cell content

Measurement of GSH/GSSG content was performed using the GSH/GSSG-Glo™ Assay (Promega; Madison, WI, USA) according to manufacturer’s instruction. Briefly 5* 10^5^ CD4^+^ T-cells were lysed and incubated with a specific probe emitting luciferin in glutathione-dependent reaction. For detection of GSSG content, N-ethylmaleimide was included to block the GSH-luciferin reaction. After addition of luciferase, a luminescent signal was developed and acquired with a Infinite 200 PRO (Tecan, Männedorf, Switzerland) multimode plate reader. A standard curve was used to calculate the total glutathione and GSSG content while the GSH/GSSG ratio was calculated as: [µM total glutathione – (µM GSSG)/µM GSSG]

### Flow Cytometry

For surface staining, 500*10^5^ cells were pelleted and fixed in PBS + 4% paraformaldehyde (PFA). Cells were then washed twice with cold FACS buffer (PBS + 2% FCS) and incubated in the dark at 4° C with an anti-TfR1 (CD71) antibody (clone OKT9; Thermo Fisher Scientific, Waltham, MA, USA). Cells were then washed with PBS and resuspended in FACS buffer. For intracellular staining, the two initial washes were performed with PBS containing 0.5% Triton x-100 and cells were stained for 30 min with an α-p24 (gag) antibody (Coulter Clone KC57-RD1; Beckman Coulter). Data were acquired with a BD FACSVerse flow cytometer and analyzed using the FlowJo package (FlowJo LLC, Ashland, Oregon, USA v7.6.5).

### RNA and DNA extraction and quantification

Total RNA and genomic DNA were extracted using, respectively, the InviTrap® Spin Universal RNA Mini Kit (Stratec Biomedical, Germany) and the DNeasy Blood & Tissue Kits (Qiagen, Germany) according to manufacturers’ instruction. Nucleic acids were quantified using a P-class P 300 NanoPhotometer (Implen GmbH, Munich, Germany).

### Reverse transcription and quantitative polymerase chain reaction (qPCR)

Total RNA was retrotranscribed to cDNA using the SuperScript III Reverse Transcriptase kit (Thermo Fisher Scientific, Waltham, MA, USA) according to manufacturer’s instruction. Briefly, 500ng of RNA were mixed with 1μL random primers (3μg/μL) and 1μL dNTPs (10mM) and incubated at 65° C per 5 min for a predenaturation step. For primer extension, 6 μL of 5X first strand buffer, 1μL DTT (0.1 M) and 0,5 μL of protector rnase inhibitor (40 U/μL, Hoffmann-La Roche, Basel, Switzerland) were added and the mix incubated at 37° C per 2 min. Finally, 1μL of MMLV RT enzyme (200 U/μL) was added and samples reverse transcribed with the following conditions: 10 min at 25° C followed by 50 min at 37° C – and 15 min at 70° C.

For qPCR, a master mix was prepared containing, per each sample, 10μL of Taq PCR Iq supermix (Bio-Rad Laboratories, Hercules, CA, USA), 1μL of primer/probe set (Applied Biosystems, Thermo Fisher) and 8μL H2O. Commercially available primer-probe mixes (Single Tube TaqMan Gene Expression Assays; Thermo Fisher Scientific, Waltham, MA, USA) were used for thioredoxin (Trx; hs01555214) used, thioredoxin reductase (TrxR1; hs00917067), glucose-6-phosphate dehydrogenase (G6PD hs00166169), heme oxygenase 1 (HMOX-1; hs01110250), glutamate-cysteine ligase (GCLC; hs00155249), NADPH dehydrogenase quinone1 (NQO1; hs02512143), Promyelocytic leukemia protein (PML; hs00231241). For U1A (gag) the following primers and probe were used: forward ACATCAAGCAGCCATGCAAAA (position 543), reverse CAGAATGGGATAGATTGCATCCA (position 629), probe AAGAGACCATCAATGAGGAA (position 605). For all qPCR reactions, 1μL of cDNA was added to the mix and qPCR reaction was performed using a CFX96/C1000 Touch qPCR system with PCR program: Polymerase activation/DNA denaturation 98°C 3 min, followed by 45 cycles of Denaturation at 98° C for 10 s; Annealing/Extension at 60° C for 40 s. Final extension was performed at 65° C for 30 s, followed by slow cool down to 4° C −0.5° C/s.

### Integrated HIV DNA measurement

The integrated HIV-1 DNA content was assessed using a nested Alu-LTR PCR assay as previously described(Tan et al. 2006). Briefly, in the first round of PCR, Alu-LTR fragments were amplified starting from 250ng of genomic DNA. For the second round, products of the first PCR were diluted 1:50 in H_2_O and LTR-LTR fragments amplified in a qPCR. A housekeeping gene, *i.e.* lamin B2, B13 region, was run in parallel in the second round of PCR using 10ng of genomic DNA and used to normalize the data as described in the statistical analyses section.

### MTT assay

Viability upon treatment with the drugs employed was measured through the CellTiter 96® Non-Radioactive Cell Proliferation Assay (MTT) assay (Promega; Madison, WI, USA) similarly to (Shytaj et al. 2015). Briefly, 300*10^5^ cells were resuspended in 100 μL RPMI + 10% FCS and transferred to a 96-well plate. To each well was added the MTT solution (15 μL) and, after 2-4 hr, the reaction was stopped by the addition of 100 μL of the Solubilization/Stop Solution. Absorbance values at 570 nm were acquired with an Infinite 200 PRO (Tecan, Männedorf, Switzerland) multimode plate reader. Reactions were conducted in triplicate and the averages of the triplicates were normalized over the matched untreated controls and expressed as percentage.

### Immunofluorescence and Immuno HIV-1 DNA FISH

Approximately 3×10^5^ uninfected or HIV-1 infected CD4^+^ T cells were plated on the PEI coated coverslips placed into a 24-well plate for 1 hr at 37° C. Cells were treated with 0.3x PBS to induce hypotonic shock and fixed in 4% PFA in PBS for 10 min. Coverslips were extensively washed with PBS and cells were permeabilized in 0.5% triton X-100/PBS for 10 min. After three additional washings with PBS-T (0.1% tween-20), coverslips were blocked with 4% BSA/PBS for 45 min at RT and primary antibody anti-rabbit lamin B1 (abcam ab16048), mab414 (1:500) (Abcam 24609), anti-mouse PML (Santa Cruz (PG-M3): sc-966 or Bethyl Lab A301-167A), (1:500 in 1% BSA/PBS) or TfR1 (1:200) (Abcam ab8598) were incubated overnight at 4° C. Following three washings with PBS-T, fluorophore-coupled secondary antibodies (anti-mouse, coupled to Alexa 488, anti-rabbit, coupled to Alexa 568 or Alexa 647, diluted 1:1000 in 1% BSA/PBS) were incubated for 1h at RT, extensively washed and postfixed with EGS in PBS. Coverslips were washed three times with PBS-T and incubated in 0.5% triton X-100/0.5% saponin/PBS for 10 min. After three washings with PBS-T, coverslips were treated with 0.1 M HCl for 10 min, washed three times with PBS-T and additionally permeabilized step in 0.5% triton X-100/0.5% saponin/PBS for 10 min. After extensive PBS-T washings, RNA digestion was performed using RNAseA (100ug/ml) for 45 min at 37° C. Coverslips were equilibrated for 5 min in 2x SSC and put in hybridization solution over night at 4°C. HIV-1 FISH probes were generated by labeling HIV-1 DNA (pHXB2) plasmid. Biotin-dUTP nucleotide mix containing 0.25 mM dATP, 0.25 mM dCTP, 0.25 mM dGTP, 0.17 mM dTTP and 0.08 mM biotin-16-dUTP in H_2_O was prepared. 3 μg of pHXB2 were diluted with H_2_O in a final volume of 12μl, and 4 μl of each nucleotide mix and Nick translation mix (Hoffmann-La Roche, Basel, Switzerland) were added. Labeling was performed at 15° C for 5 h. The probe was precipitated in 100% ethanol with sodium acetate overnight and resuspended in 2x SSC/10% dextran sulfate/50% formamide, denatured and stored at −20° C until use. Probe hybridization was performed with 2μl of HIV-1 probe in 6 μl reaction with 2x SSC/10% dextran sulfate/50% formamide, denatured at 95° C for 5 min and then kept on ice for 1 min. After spotting the probe on a glass slide in a metal chamber, and put at 80° C for 8 min for denaturation and subsequent incubation at 37°C for 40 to 65 h in a water bath to allow hybridization of the probe. Probe detection was carried out by washing the coverslips with 2x SSC and 0.5x SSC at 37 and 65° C, respectively; 1hr blocking in TSA blocking buffer (TNB) and detection with streptavidin-HRP in TNB for 40 min at 37° C. Coverslips were then washed with TNT wash buffer at RT, before incubation with Fluorescein Plus amplification reagent (1:1500 in TSA Plus amplification diluent, part of TSA Plus Fluorescein kit) for 5 min at RT. Final five washings with TNT buffer and nuclear counterstaining was performed using 1:10000. Hoechst 33342 in PBS followed by two washings in PBS and mounting the coverslips with mowiol.

### Single molecule HIV-1 RNA FISH coupled to immunofluorescence

A custom made Stellaris HIV-1 RNA FISH probe Fluor Red 610, dissolved to 12.5 μM in TE buffer was used for the RNA FISH experiments. Approximately 5×10^5^ HIV-1 infected PBMCs were washed twice in RNA grade PBS, and fixed in 3.7% Formaldehyde (FA) solution in PBS for 10 min. After extensive washings with PBS, cells were permeabilized in suspension in 70% ice-cold ethanol for at least 1hr. Cells were then adhered to PEI coated slides for 30 min at RT and washed with PBS RNA grade for 3 times 3 min at RT. IF for Nrf2 (mab3925, 1:250; R&D systems, Minneapolis, MN) or PML (sc/966x, 1:500; Santa Cruz Biotechnology, Dallas, TX, USA) for 1-2 hr at RT. After PBS washings, secondary antibody anti-mouse, coupled to Alexa 488 (1:1000 dilution) was incubated 1 hr at RT light protected.

After extensive PBS washings, cells were post-fixed with 3.7 % FA in RNA grade PBS at room temperature for 10 min, washed with PBS and permeabilized with 70% ice cold ethanol for 30 min. Cells were washed/equilibrated in buffer A (Stellaris RNA FISH Wash Buffer A (Biosearch Tech. Cat# SMF-WA1-60) with 10% formamide in water. 1-3 μL of HIV-1 RNA probe diluted in hybridization buffer (Stellaris RNA FISH Hybridization Buffer (Biosearch Tech. Cat# SMF-HB1-10)) and 10% formamide were spotted on coverslips and incubate overnight in humid conditions at 37C. Coverslips were subsequently washed with buffer A four times at 37° C, one time at RT, and nuclear counterstaining was performed using 1:10000 Hoechst 33342 in buffer A followed by two washings in buffer B (Stellaris RNA FISH Wash Buffer B (Biosearch Tech. Cat# SMF-WB1-20) and mounting the coverslips with mowiol.

### Stimulated emission depletion (STED) microscopy

STED imaging was performed with a λ=775 nm STED system (Abberior Instruments GmbH, Göttingen, Germany), containing an easy 3D optics module (Abberior Instruments) and the =640 nm excitation laser line using a 100x Olympus UPlanSApo (NA 1.4) oil immersion objective. Images were acquired using the 590 and 640 nm excitation laser lines.

Additional confocal images of eGFP signals excited at 488 nm were acquired. Deconvolution of STED images was performed in Imspector software (Abberior Instruments) via the linear deconvolution tool.

Images were further processed in Fiji-Imagej (Schindelin et al. 2012) by using the Gaussian-Blur Filter and setting the sigma radius to 1 to further increase signal/noise ratio. Different analyzed excitation channels from the same sample were also merged into one image by using this software.

### Confocal microscopy

Data were acquired using a Leica TCS SP8 confocal microscope (Leica Microsystems GmbH, Wetzlar, Germany) with a 63x objective immersed in oil. A distance (z-step) of 500 nm was used to acquire three dimensional stacks with zoom set at 1x or 3x.

Image analysis was performed with Fiji-Imagej (Schindelin et al. 2012) using the following macros to perform automatic segmentation and counting of the number of cells and PML bodies:

Segmentation:

~~~
//run(“Brightness/Contrast…”);
setMinAndMax(0, 150);
run(“Gaussian Blur…”, “sigma=2 stack”);
//run(“Threshold…”); setAutoThreshold(“RenyiEntropy dark”);
setThreshold(40, 255);
//setThreshold(40, 255);
setOption(“BlackBackground”, true);
run(“Convert to Mask”, “method=RenyiEntropy background=Dark black”);
roiManager(“Show None”);
roiManager(“Show All”);
~~~

count:

~~~
run(“Duplicate…”, “duplicate”);
run(“3D Objects Counter”, “threshold=1 slice=16 min.=1 max.=356796 objects statistics summary”);
~~~

### 3D FISH PML analysis algorithm

We developed an algorithm, implemented in MATLAB, in order to automatically analyse 3D RNA FISH and PML proteins in fluorescence cell image z-stacks. The algorithm is derived from Gregoretti *et al.* (manuscript submitted) with some adaptations. It performs the 2D segmentation of cell nuclei and the detection of FISH and PML spots for each slice of the stack followed by the 3D reconstruction and identification of nuclei and spots. It then counts and measures the 3D FISH and PML spots and calculate the total intensity value of FISH spots. We recognized as infected those cells with HIV-1 RNA FISH signals satisfying these constraints: 1) at least one signal with size greater than 200 voxels; 2) total intensity of FISH signals greater than 15000. The algorithm can be sketched as follows:

for each slice n of the stack

~~~
     I_vol_FISH_ = I_FISH,n_(:,:)
     nuclei_n_ = ***nuclei_seg***(I_DAPI,n_) %performs 2D nuclei segmentation
     nuclei_vol(:,:,n) = nuclei_n_(:,:)
     fish_n_ = ***detect_spot***(I_FISH,n_) %performs 2D FISH spot detection
     fish_vol(:,:,n) = fish_n_(:,:)
     pml_n_ = ***detect_spot***(I_PML,n_) %performs 2D PML spot detection
     pml_vol(:,:,n) = pml_n_(:,:)
endfor
nuclei_CC = **bwconncomp**(nuclei_vol)
nuclei_L = **labelmatrix**(nuclei_CC)
compute volume for each nucleus object in nuclei_CC
exclude nuclei whose volume is less than 10% of mean volumes
{NCL}_M_ <-identified 3D nuclei pixel data
for each nucleus m in {NCL}_M_
    NCL_m_.FISH = {NCL}_M._* fish_vol %3D positions of detected FISH spots within NCL_m_
    NCL_m_.FISH = **bwareaopen**(NCL_m_.FISH,17,6)
    FISH_CC_m_ = **bwconncomp**(NCL_m_.FISH)
    compute volume for each FISH spot object in FISH_CC_m_
    exclude FISH spots whose volume is below 100 voxels
    NCL_m_.FISH_PICK_ <-final FISH spot objects
    count final FISH spot objects
    NCL_m_.Intensity_FISH_ = I_vol_FISH._* NCL_m_.FISH_PICK_ %intensity values of FISH spots
    NCL_m_.TotIntensity_FISH_ = sum(sum(sum(NCL_m_.Intensity_FISH_))); %total intensity value of FISH spots
    NCL_m_ PML = {NCL}_M._* pml_vol %3D positions of detected PML spots within NCL_m_
    NCL_m_.PML = **bwareaopen**(NCL_m_.PML,17,6)
    PML_CC_m_ = **bwconncomp**(NCL_m_.PML)
    compute volume for each PML spot object in PML_CC_m_
    exclude PML spots object whose volume is below 0.15 micron cubes
    NCL_m_.PML_PICK_ <-final PML spot objects
    count final PML spot objects
endfor
~~~

*I*_*DAPI*_, *I*_*FISH*_ and *I*_*PML*_ are the stack images with DAPI, RNA FISH and PLM staining respectively. The function *nuclei_seg* performs a partition of cell image in nuclei regions and background implementing a region based segmentation algorithm(Chan et al. 2006; Goldstein et al. 2009; Antonelli & De Simone 2018). Output image *nuclei*_*n*_ highlights foreground objects (nucleus regions). The function *detect_spot* has four major steps. It first filters the input image *I*_*FISH,n*_ (*I*_*PML.n*_) applying the Laplacian of Gaussian (LoG) operator (*fspecial* MATLAB function) of size 9 and standard deviation 7. This enhances the signal in the areas where objects are present. Then the function applies the h-dome transformation (Vincent 1993) that extracts bright structures by cutting off the intensity of height h from the top, around local intensity maxima. We used h=0.5 with a neighborhood size of 15×15. We decided to not use a global operator after having observed that a FISH spot (PML spot) in one part of the image could be lighter or darker than the background in another part. This is due to the facts that spots have inhomogeneous intensity distribution over the image and that the image may have an uneven background. In the third step, the function performs a thresholding on h-domes image that excludesGold pixels whose intensity values are below a threshold. The FISH and PML threshold values are defined as 1.96 and 1 standard deviations above the mean of domes intensity values, respectively. We therefore assumed that spot areas have significant intensity disparity with respect to other bright areas present in cell nucleus. Lastly, the function applies a thresholding operation based on the surface areas of the spots, in order to discard too small objects which are probably just noise. It filters out spots smaller than a surface area of 8. *detect_spot* produces an accurate set of FISH and PML spots.

*fish_vol, pml_vol* and *nuclei_vol* are 3D arrays that contain the positions of the detected FISH and PML spots and nuclei from all slices.

3D reconstructions of nuclei are obtained through the connected components algorithm (*bwconncomp* MATLAB function, using a connectivity of 6). 3D nuclei are then labeled by applying the *labelmatrix* MATLAB function so they can easily separated each from the others.

The algorithm computes the volume of each 3D reconstruction, discarding, as noise, objects whose volume is less than 10% of mean volumes.

The algorithm uses the *bwareaopen* function in order to discard too small detected spot objects which are probably just noise. 3D reconstructions of spots are obtained through the connected components algorithm (*bwconncomp* MATLAB function, using a connectivity of 6). Then a threshold operation is performed on the 3D spots to obtain the more significant ones: it keeps in all the PML spots whose volume is above 0.15 micron cubes and all the FISH spots whose volume is above 100 voxels.

### Biochemical fractionation, SDS-PAGE, and Western Blot

Cytoplasmic and nuclear cell fractions were prepared from 10 to 20×10^6^ cells. Briefly, cell pellets were resuspended in cytoplasmic buffer: 10mM Tris pH 7.9; 3mM CaCl_2_, 2mM MgCl_2_, 0.1 mM EDTA, 0.34 M sucrose, 1mM DTT supplemented with protease inhibitors (Roche). After swelling in cytoplasmic buffer on ice for 5 min, Triton X-100 was added to the final concentration of 0.1%. Cytoplasm was extracted after 10 min incubation and 10 min centrifugation at 3000 rpm at 4C. Nuclei were washed in the same buffer without the detergent, and lysed in nuclei lysis buffer: 20 mM Hepes pH 7.9; 2.5 mM MgCl2; 0.5mM EDTA; 150mM KCl; 10% glycerol; 0.5% NP 40 plus protease inhibitors (add fresh) on ice for 10 min. Nucleic acids were removed with 250 U of Benzonase (Roche) for 1 hr at 4° C. Nuclear soluble fraction was extracted by addition of high salt buffer to achieve 250 mM KCl for 1 hr at 4°C rotating, followed by 45 min centrifugation at maximum speed at 4°C. Alternatively, cytoplasmatic and nuclear cell fractions were isolated using the REAP method, according to the published protocol (Suzuki et al. 2010).

For SDS-Page experiments 1-5* 10^6^ cells were harvested and homogenized in lysis buffer (20 mM Tris-HCl, pH 7.4, 1 mM EDTA, 150 mM NaCl, 0.5% Nonidet P-40, 0.1% SDS, 0.5% sodium deoxycholate supplemented with protease inhibitors (Hoffmann-La Roche, Basel, Switzerland) for 10 min at 4° C. For further homogenization, lysates were sonicated for 5 min using a Bioruptor® Plus sonication device (Diagenode, Liège, Belgium). Protein concentration was assayed using the Micro bca protein assay kit (Thermo Fisher Scientific, Waltham, MA, USA) or Bradford method (Bio-Rad Laboratories, Hercules, CA, USA) and acquired through a Implen NanoPhotometer® Pearl (Implen GmbH, Munich, Germany). Equal amounts of total cellular proteins (10 or 25 μg), were loaded and run on a precast NuPAGE Bis-Tris 4-12% (Thermo Fisher Scientific, Waltham, MA, USA) SDS-PAGE at 120V. Proteins were then transferred onto a nitrocellulose membrane (GE Healthcare, Little Chalfont, UK) for 1 hr using a Trans–Blot device for semi-dry transfer (Bio-Rad Laboratories, Hercules, CA, USA). Membranes were then blocked for 1 hr at RT with 5% skim milk in 0,1 % PBS-Tween and incubated overnight at 4o with the following primary antibodies diluted in 5% milk: α-beta-actin (1:1000), (Sigma Aldrich, Saint Louis, MI, USA), α-Trx (1:500; sc-58440), α-NQO1 (1:500, sc-32793) (Santa Cruz Biotechnology, Dallas, TX, USA), α-TrxR1 (1:500, MAB7428) (R&D systems, Minneapolis, MN), α-HMOX-1 (1:500, ab13248), FTH-1 (1:100, ab75972), α SLC40A1 (1:500, ab78066) (Abcam, Cambridge, UK) α-PML (1:500) (A301-167A Bethyl Laboratories, Montgomery, TX, USA). Membranes were then washed three times with 0,1 % PBS-Tween and incubated for 1 hr with a horseradish-conjugated secondary antibody (α-mouse or anti rabbit, West Grove, PA, USA). Proteins were visualized with the Amersham ECL Prime kit (GE Healthcare, Little Chalfont, UK). For membrane reprobing, stripping buffer was used (2% SDS, 8% Upper Tris buffer and 110 mM ß-mercaptoethanol) for 15 min at 65° C. After extensive washing with 0,1 % PBS-Tween, membrane was blocked and reincubated with desired antibodies.

### Statistical analysis

qPCR and Alu PCR values were normalized using the 2(-ΔΔ C(T)) method (Livak & Schmittgen 2001). Comparisons between two groups were performed by unpaired *t*-test. Three or more groups were compared by either one way or two way ANOVA followed by the appropriate post-test for multiple comparisons.

## Supplementary figures

**Figure S1**. **Influence of HIV-1 replication on protein expression of antioxidant genes.** Proteomic analysis of the relative expression of antioxidant genes in primary CD4^+^ T-cells infected *in vitro* with HIV-1 or mock infected. N of donors = 3

**Figure S2. Individual patterns of antioxidant responses and virological features of the HIV-1 mutants employed in the study.** (A) Single donor dynamics of the data shown in Figure 3C. (B, C) relative *gag* mRNA (B) and integrated viral DNA (C) expression levels of HIV-1_NL4-3_ viruses bearing mutations in *nef, tat* or *env* as compared to *wt*. RNA and viral DNA levels were measured by qPCR or Alu-HIV PCR respectively and expressed as Log_2_ fold change expression in infected over mock infected (panel E) or mutant over *wt* HIV-1 (B, C) using the 2-ΔΔCTmethod (Livak & Schmittgen 2001).

**Figure S3. Regulation of iron import and cell viability upon HIV-1 infection or treatment with FeCl_3_ 6H_2_O** (A) Proteomic analysis of different time points in primary CD4^+^ T-cells infected *in vitro* with HIV-1 or mock infected. The gene sets used for the analyses are *GO cellular ion iron homeostasis* (46 genes, GO:0006879; left panel) and *GO iron ion import* (12 genes, GO:0097286; right panel). (B) relative TfR1 expression over time in HIV-1 infected vs mock infected CD4^+^ T-cells as measured by flow cytometry. (C) representative image of TfR1 expression in CD4^+^ T-cells resting or activated for 72 hr with α-CD3/CD28 beads. (D) relative viability of CD4^+^ T-cells incubated for 48 hr with various concentrations of the iron donor FeCl_3_ 6H_2_O. Viability was measured using the MTT assay. Absorbance values were normalized over untreated controls and expressed as percentage (mean±SEM, n=3). Data were analyzed by one-way ANOVA followed by Dunnet’s multiple comparisons test.

**Figure S4**. **Impact of oxidative stress and HIV-1 replication on PML stability** (A,B) Nrf2 subcellular localization (A) and PML expression (B) in CD4^+^ T-cells left untreated or treated for 30 min with 100μM H_2_O_2_. After washing away H_2_O_2_ cells were cultured for 24 hours and harvested for biochemical fractionation (A) and western blot (A,B). (C,D) Representative 3D reconstruction (C) and quantification (D) of the number of PML NBs in HIV-1 RNA^+^ vs HIV-1 RNA^-^ CD4^+^ T-cells at 7dpi. Infected cells were identified by RNA FISH and PML NBs were stained by IF. The algorithm used for 3D cell reconstruction and PML counting is detailed in the materials and methods section. Data were analyzed by two-tailed unpaired *t*-test and are expressed as mean±SEM; n= number of cells from 3 donors.

**Figure S5 Effect of pharmacologic PML depletion on HIV-1 replication and integration.** (A-C) PML expression in CD4^+^ T-cells left untreated or treated for 48 hr with 5μM As_2_O_3_ as measured by IF (A,B) or western blot (C). n= number of cells from 3 donors; mean±SEM. Data were analyzed by unpaired *t*-test. (D,E) quantification (D) and representative image (E) of the effect of PML depletion through As_2_O_3_ on HIV-1 transcription/production and integration/latency. CD4^+^ T-cells were incubated with 5μM As_2_O_3_ or left untreated. After 4-6 hr treatment cells were infected with *wt* HIV-1_NL4-3_ or the dual color replication-deficient OGH HIV-1 virus (Vranckx et al. 2016; Calvanese et al. 2013). At day 1 and/or 3 post-infection cells were assayed for: HIV-1 mRNA or percentage of productively infected cells (D, left), integrated HIV-1 DNA or percentage of latently infected cells (D, middle) and ratio between viral production/integration (D, right). Viral production was measured by qPCR (*wt* HIV-1_NL4-3_) or as percentage of GFP^+^ (productively infected) cells (OGH HIV-1). Viral integration was measured by nested Alu-LTR (*wt* HIV-1_NL4-3_) or as percentage of Kusabira Orange^+^ (latently infected) cells (OGH). Data are expressed as mean±SEM fold change expression over the corresponding control and analyzed by one-way ANOVA followed by Turkey’s post-test. For *wt* HIV-1_NL4-3_ (n=3) for OGH HIV-1 (n=4).

**Figure S6**: **Effect of iron chelation on TfR1 expression, HIV-1 replication and viability of CD4^+^ T-cells.** (A) representative image of TfR1 expression in activated CD4^+^ T-cells left untreated or treated with 50μM deferiprone (L1). (B) relative viability of CD4^+^ T-cells incubated for 72 hr with different concentrations of the iron chelators deferoxamine (DFOA, left) and L1 (right). Arrows indicate the concentrations used for further experiments. Viability was measured using the MTT assay. Absorbance values were normalized over untreated controls and expressed as percentage (mean±SEM, n=3). (C) percentage of p24^+^ gag CD4^+^ T-cells left untreated or treated with 50μM L1. (D) time course of the relative viability of HIV-1- or mock infected CD4^+^ T-cells left untreated or treated with 50μM L1. Viable cells were identified by flow cytometry through FCS/SSC gating (mean±SEM, n=3). Data were analyzed by one way ANOVA followed by Dunnet’s post-test (A) or by two way ANOVA followed by Sidak’s post-test (C,D). **** *P*<0,0001; *** *P*<0,001; ** *P*<0,01; * *P*<0,5.

## Acknowledgments

The authors thank Dr. Oliver Fackler and Dr. Hans Georg Kräusslich for kindly providing the HIV-1_NL4-3_ Δ*nef* and Δ*tat* vectors, and Dr. Eric Verdin for dual-color HIV reporter vector. The authors acknowledge the Infectious Diseases Imaging Platform (IDIP) and thank the platform coordinator Dr. Vibor Laketa for valuable advice. This work was supported by German Center for Infection Research (DZIF). ILS acknowledges the post-doctoral fellowship and funding provided by the Humboldt Foundation.

## Author Contributions

I.L.S, B.L, A.S and M.L conceived the project. I.L.S, B.L, B.G, G.S, A.S and M.L designed the experiments. I.L.S and B.L. performed *in-vitro* experiments and analyzed *in-vitro* data. M.F., S.B., B.J.M. and B.S performed bioinformatic analysis. C.E performed the molecular modelling. G.F, A.L and O.G performed 3D cell reconstructions. M.S. analyzed STED microscopy data. I.L.S, B.L. A.S. and M.L. wrote the manuscript.

## Bibliography

Antonelli, L. & De Simone, V., 2018. Comparison of minimization methods for nonsmooth image segmentation. Communications in Applied and Industrial Mathematics, 9(1), pp.68–86.

Bejarano, D.A. et al., 2019. HIV-1 nuclear import in macrophages is regulated by CPSF6-capsid interactions at the Nuclear Pore Complex. eLife, 8. Available at: http://dx.doi.org/10.7554/eLife.41800.

Benhar, M. et al., 2016. Dual targeting of the thioredoxin and glutathione systems in cancer and HIV. The Journal of clinical investigation, 126(5), pp.1630–1639.

Bhaskar, A. et al., 2015. Measuring glutathione redox potential of HIV-1-infected macrophages. The Journal of biological chemistry, 290(2), pp.1020–1038.

Bosque, A. & Planelles, V., 2009. Induction of HIV-1 latency and reactivation in primary memory CD4+ T cells. Blood, 113(1), pp.58–65.

Calvanese, V. et al., 2013. Dual-color HIV reporters trace a population of latently infected cells and enable their purification. Virology, 446(1-2), pp.283–292.

Chan, T.F., Esedoglu, S. & Nikolova, M., 2006. Algorithms for Finding Global Minimizers of Image Segmentation and Denoising Models. SIAM journal on applied mathematics, 66(5), pp.1632–1648.

Chavez, L., Calvanese, V. & Verdin, E., 2015. HIV Latency Is Established Directly and Early in Both Resting and Activated Primary CD4 T Cells. PLoS pathogens, 11(6), p.e1004955.

Chirullo, B. et al., 2013. A candidate anti-HIV reservoir compound, auranofin, exerts a selective “anti-memory” effect by exploiting the baseline oxidative status of lymphocytes. Cell death & disease, 4, p.e944.

Chomont, N. et al., 2009. HIV reservoir size and persistence are driven by T cell survival and homeostatic proliferation. Nature medicine, 15(8), pp.893–900.

Chun, T.W. et al., 1998. Early establishment of a pool of latently infected, resting CD4(+) T cells during primary HIV-1 infection. Proceedings of the National Academy of Sciences of the United States of America, 95(15), pp.8869–8873.

Chun, T.W. et al., 1999. Re-emergence of HIV after stopping therapy. Nature, 401(6756), pp.874–875.

Ciciliano, J.C. et al., 2015. Resolving the multifaceted mechanisms of the ferric chloride thrombosis model using an interdisciplinary microfluidic approach. Blood, 126(6), pp.817–824.

Condemine, W. et al., 2006. Characterization of endogenous human promyelocytic leukemia isoforms. Cancer research, 66(12), pp.6192–6198.

Dobin, A. et al., 2013. STAR: ultrafast universal RNA-seq aligner. Bioinformatics, 29(1), pp.15–21.

Drakesmith, H. & Prentice, A., 2008. Viral infection and iron metabolism. Nature reviews. Microbiology, 6(7), pp.541–552.

Dutrieux, J. et al., 2015. PML/TRIM19-Dependent Inhibition of Retroviral Reverse-Transcription by Daxx. PLoS pathogens, 11(11), p.e1005280.

Evans, D.T. & Silvestri, G., 2013. Nonhuman primate models in AIDS research. Current opinion in HIV and AIDS, 8(4), pp.255–261.

Furuya, A.K.M. et al., 2016. Sulforaphane Inhibits HIV Infection of Macrophages through Nrf2. PLoS pathogens, 12(4), p.e1005581.

Georgiou, N.A. et al., 2002. Human immunodeficiency virus type 1 replication inhibition by the bidentate iron chelators CP502 and CP511 is caused by proliferation inhibition and the onset of apoptosis. European journal of clinical investigation, 32 Suppl 1, pp.91–96.

Georgiou, N.A. et al., 2000. Inhibition of human immunodeficiency virus type 1 replication in human mononuclear blood cells by the iron chelators deferoxamine, deferiprone, and bleomycin. The Journal of infectious diseases, 181(2), pp.484–490.

Gill, A.J. et al., 2014. Heme oxygenase-1 deficiency accompanies neuropathogenesis of HIV-associated neurocognitive disorders. The Journal of clinical investigation, 124(10), pp.4459–4472.

Giustarini, D. et al., 2013. Analysis of GSH and GSSG after derivatization with N-ethylmaleimide. Nature protocols, 8(9), pp.1660–1669.

Goldstein, T., Bresson, X. & Osher, S., 2009. Geometric Applications of the Split Bregman Method: Segmentation and Surface Reconstruction. Journal of scientific computing, 45(1-3), pp.272–293.

Gorrini, C., Harris, I.S. & Mak, T.W., 2013. Modulation of oxidative stress as an anticancer strategy. Nature reviews. Drug discovery, 12(12), pp.931–947.

Gozzelino, R., Jeney, V. & Soares, M.P., 2010. Mechanisms of Cell Protection by Heme Oxygenase-1. Annual review of pharmacology and toxicology, 50(1), pp.323–354.

Guo, S. et al., 2014. Control of antioxidative response by the tumor suppressor protein PML through regulating Nrf2 activity. Molecular biology of the cell, 25(16), pp.2485–2498.

Gu, Y. et al., 2001. HIV Tat Activates c-Jun Amino-terminal Kinase through an Oxidant-Dependent Mechanism. Virology, 286(1), pp.62–71.

Hands, K.J. et al., 2014. PML isoforms in response to arsenic: high-resolution analysis of PML body structure and degradation. Journal of cell science, 127(Pt 2), pp.365–375.

Hiener, B. et al., 2017. Identification of Genetically Intact HIV-1 Proviruses in Specific CD4 T Cells from Effectively Treated Participants. Cell reports, 21(3), pp.813–822.

Hughes, C.S. et al., 2019. Single-pot, solid-phase-enhanced sample preparation for proteomics experiments. Nature protocols, 14(1), pp.68–85.

Isaguliants, M. et al., 2013. Oxidative stress induced by HIV-1 reverse transcriptase modulates the enzyme’s performance in gene immunization. Human vaccines & immunotherapeutics, 9(10), pp.2111–2119.

Kahle, T. et al., 2015. TRIM19/PML Restricts HIV Infection in a Cell Type-Dependent Manner. Viruses, 8(1). Available at: http://dx.doi.org/10.3390/v8010002.

Kerins, M.J. & Ooi, A., 2018. The Roles of NRF2 in Modulating Cellular Iron Homeostasis. Antioxidants & redox signaling, 29(17), pp.1756–1773.

Klammer, H. et al., 2015. Bystander effects as manifestation of intercellular communication of DNA damage and of the cellular oxidative status. Cancer letters, 356(1), pp.58–71.

Liberzon, A. et al., 2011. Molecular signatures database (MSigDB) 3.0. Bioinformatics, 27(12), pp.1739–1740.

Livak, K.J. & Schmittgen, T.D., 2001. Analysis of relative gene expression data using real-time quantitative PCR and the 2(-Delta Delta C(T)) Method. Methods, 25(4), pp.402–408.

Love, M.I., Huber, W. & Anders, S., 2014. Moderated estimation of fold change and dispersion for RNA-seq data with DESeq2. Genome biology, 15(12), p.550.

Lusic, M. et al., 2013. Proximity to PML nuclear bodies regulates HIV-1 latency in CD4+ T cells. Cell host & microbe, 13(6), pp.665–677.

MacKenzie, E.L., Iwasaki, K. & Tsuji, Y., 2008. Intracellular iron transport and storage: from molecular mechanisms to health implications. Antioxidants & redox signaling, 10(6), pp.997–1030.

Ma, Q., 2013. Role of nrf2 in oxidative stress and toxicity. Annual review of pharmacology and toxicology, 53, pp.401–426.

Martins, L.J. et al., 2016. Modeling HIV-1 Latency in Primary T Cells Using a Replication-Competent Virus. AIDS research and human retroviruses, 32(2), pp.187–193.

McCloy, R.A. et al., 2014. Partial inhibition of Cdk1 in G2phase overrides the SAC and decouples mitotic events. Cell cycle, 13(9), pp.1400–1412.

Muckenthaler, M.U. et al., 2017. A Red Carpet for Iron Metabolism. Cell, 168(3), pp.344–361.

Niwa-Kawakita, M. et al., 2017. PML is a ROS sensor activating p53 upon oxidative stress. The Journal of experimental medicine, 214(11), pp.3197–3206.

Pace, G.W. & Leaf, C.D., 1995. The role of oxidative stress in HIV disease. Free radical biology & medicine, 19(4), pp.523–528.

Palesch, D. et al., 2018. Short-Term Pegylated Interferon α2a Treatment Does Not Significantly Reduce the Viral Reservoir of Simian Immunodeficiency Virus-Infected, Antiretroviral Therapy-Treated Rhesus Macaques. Journal of virology, 92(14). Available at: http://dx.doi.org/10.1128/jvi.00279-18.

Pietraforte, D. et al., 1994. gp120 HIV envelope glycoprotein increases the production of nitric oxide in human monocyte-derived macrophages. Journal of leukocyte biology, 55(2), pp.175–182.

Pizzato, M. et al., 2007. Dynamin 2 is required for the enhancement of HIV-1 infectivity by Nef. Proceedings of the National Academy of Sciences of the United States of America, 104(16), pp.6812–6817.

Sahin, U., Ferhi, O., et al., 2014. Oxidative stress-induced assembly of PML nuclear bodies controls sumoylation of partner proteins. The Journal of cell biology, 204(6), pp.931–945.

Sahin, U., de Thé, H. & Lallemand-Breitenbach, V., 2014. PML nuclear bodies: assembly and oxidative stress-sensitive sumoylation. Nucleus, 5(6), pp.499–507.

Savarino, A., Calosso, L., et al., 1999. Modulation of surface transferrin receptors in lymphoid cells de novo infected with human immunodeficiency virus type-1. Cell biochemistry and function, 17(1), pp.47–55.

Savarino, A., Pescarmona, G.P. & Boelaert, J.R., 1999. Iron metabolism and HIV infection: reciprocal interactions with potentially harmful consequences? Cell biochemistry and function, 17(4), pp.279–287.

Schindelin, J. et al., 2012. Fiji: an open-source platform for biological-image analysis. Nature methods, 9(7), pp.676–682.

Shytaj, I.L. et al., 2013. Investigational treatment suspension and enhanced cell-mediated immunity at rebound followed by drug-free remission of simian AIDS. Retrovirology, 10, p.71.

Shytaj, I.L. et al., 2015. Two-Year Follow-Up of Macaques Developing Intermittent Control of the Human Immunodeficiency Virus Homolog Simian Immunodeficiency Virus SIVmac251 in the Chronic Phase of Infection. Journal of virology, 89(15), pp.7521–7535.

Siliciano, R.F. & Greene, W.C., 2011. HIV latency. Cold Spring Harbor perspectives in medicine, 1(1), p.a007096.

Suzuki, K. et al., 2010. REAP: A two minute cell fractionation method. BMC research notes, 3, p.294.

Tan, W. et al., 2006. Human Immunodeficiency Virus Type 1 Incorporated with Fusion Proteins Consisting of Integrase and the Designed Polydactyl Zinc Finger Protein E2C Can Bias Integration of Viral DNA into a Predetermined Chromosomal Region in Human Cells. Journal of virology, 80(4), pp.1939–1948.

Trautz, B. et al., 2016. The Antagonism of HIV-1 Nef to SERINC5 Particle Infectivity Restriction Involves the Counteraction of Virion-Associated Pools of the Restriction Factor. Journal of virology, 90(23), pp.10915–10927.

Turelli, P. et al., 2001. Cytoplasmic recruitment of INI1 and PML on incoming HIV preintegration complexes: interference with early steps of viral replication. Molecular cell, 7(6), pp.1245–1254.

Valle-Casuso, J.C. et al., 2018. Cellular Metabolism Is a Major Determinant of HIV-1 Reservoir Seeding in CD4 T Cells and Offers an Opportunity to Tackle Infection. Cell metabolism. Available at: http://dx.doi.org/10.1016/j.cmet.2018.11.015.

Vilhardt, F. et al., 2002. The HIV-1 Nef protein and phagocyte NADPH oxidase activation. The Journal of biological chemistry, 277(44), pp.42136–42143.

Vincent, L., 1993. Morphological grayscale reconstruction in image analysis: applications and efficient algorithms. IEEE transactions on image processing: a publication of the IEEE Signal Processing Society, 2(2), pp.176–201.

Vranckx, L.S. et al., 2016. LEDGIN-mediated Inhibition of Integrase-LEDGF/p75 Interaction Reduces Reactivation of Residual Latent HIV. EBioMedicine, 8, pp.248–264.

Wang, J. & Pantopoulos, K., 2011. Regulation of cellular iron metabolism. Biochemical Journal, 434(3), pp.365–381.

Wang, P. et al., 2018. Publisher Correction: RING tetramerization is required for nuclear body biogenesis and PML sumoylation. Nature communications, 9(1), p.1841.

Williams, S.A. & Greene, W.C., 2007. Regulation of HIV-1 latency by T-cell activation. Cytokine, 39(1), pp.63–74.

Zhang, H.-S. et al., 2009. Nrf2 is involved in inhibiting Tat-induced HIV-1 long terminal repeat transactivation. Free Radical Biology and Medicine, 47(3), pp.261–268.

Zhang, X.-W. et al., 2010. Arsenic trioxide controls the fate of the PML-RARalpha oncoprotein by directly binding PML. Science, 328(5975), pp.240–243.

Zhou, X. et al., 2015. Selective Sensitization of Zinc Finger Protein Oxidation by Reactive Oxygen Species through Arsenic Binding. The Journal of biological chemistry, 290(30), pp.18361–18369.

