## Supplementary figures and images for "Alterations of redox and iron metabolism accompany development of HIV latency"

### Supplementary Figures 1-6

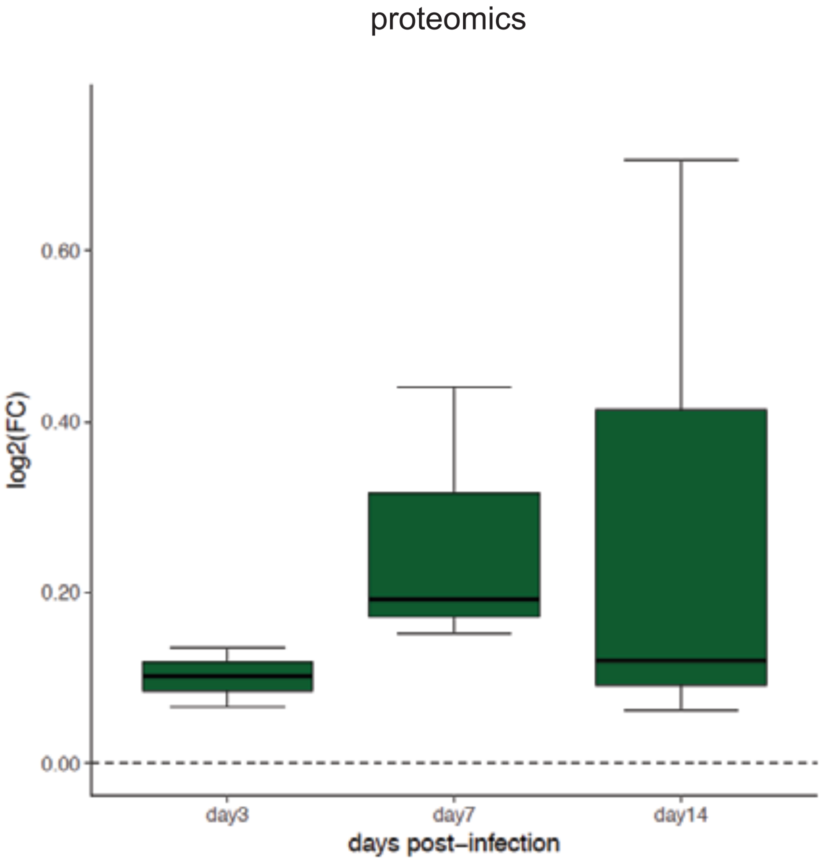

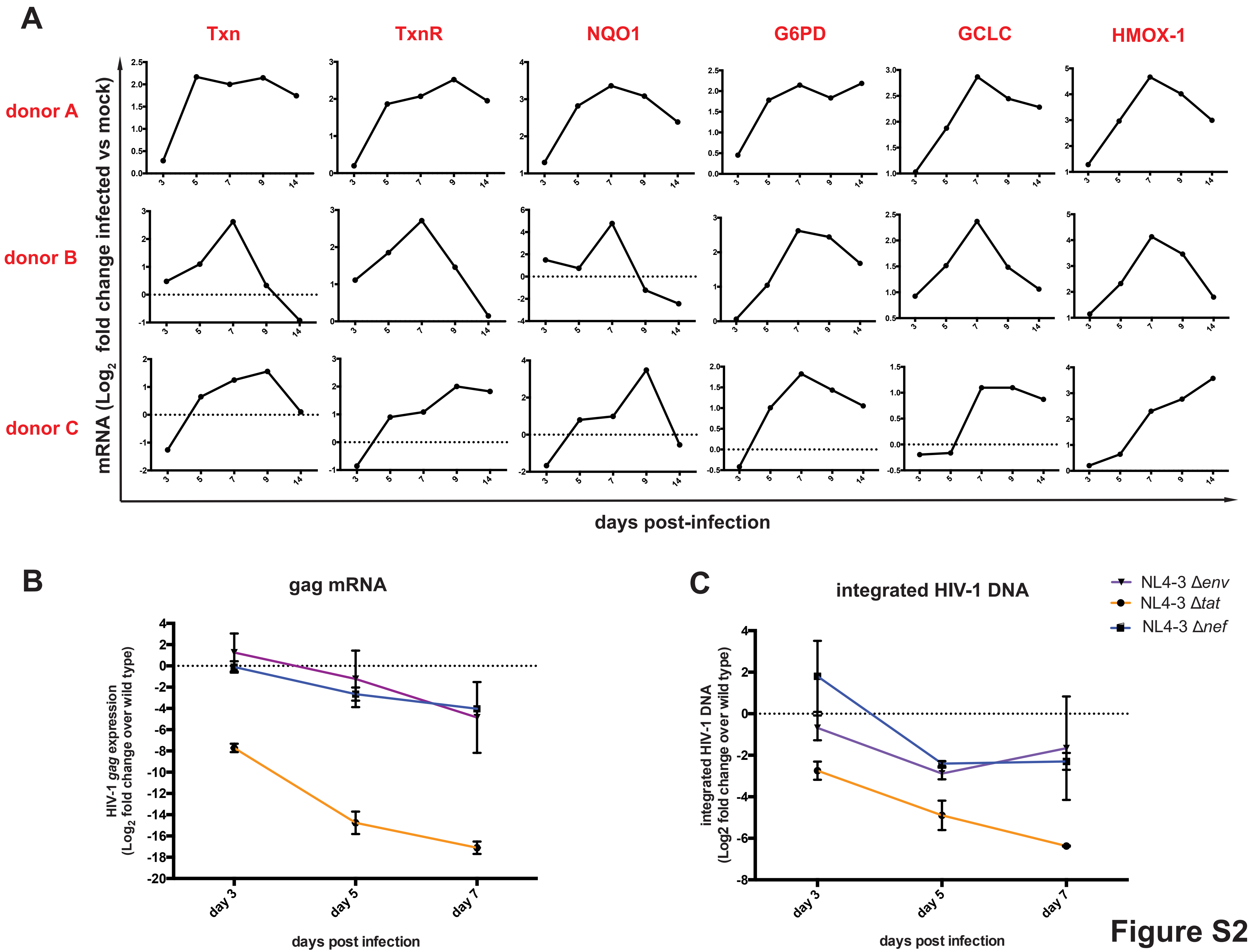

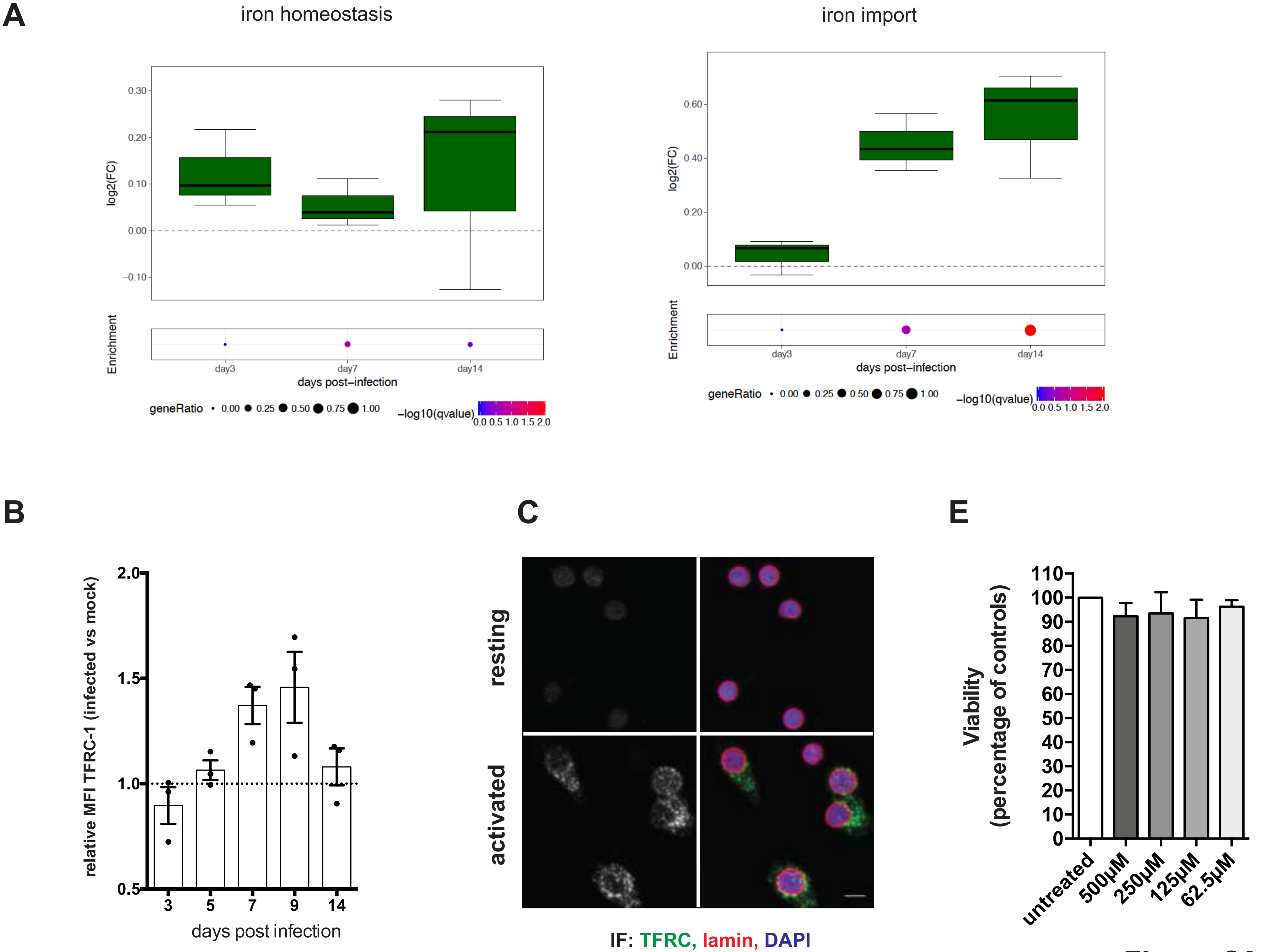

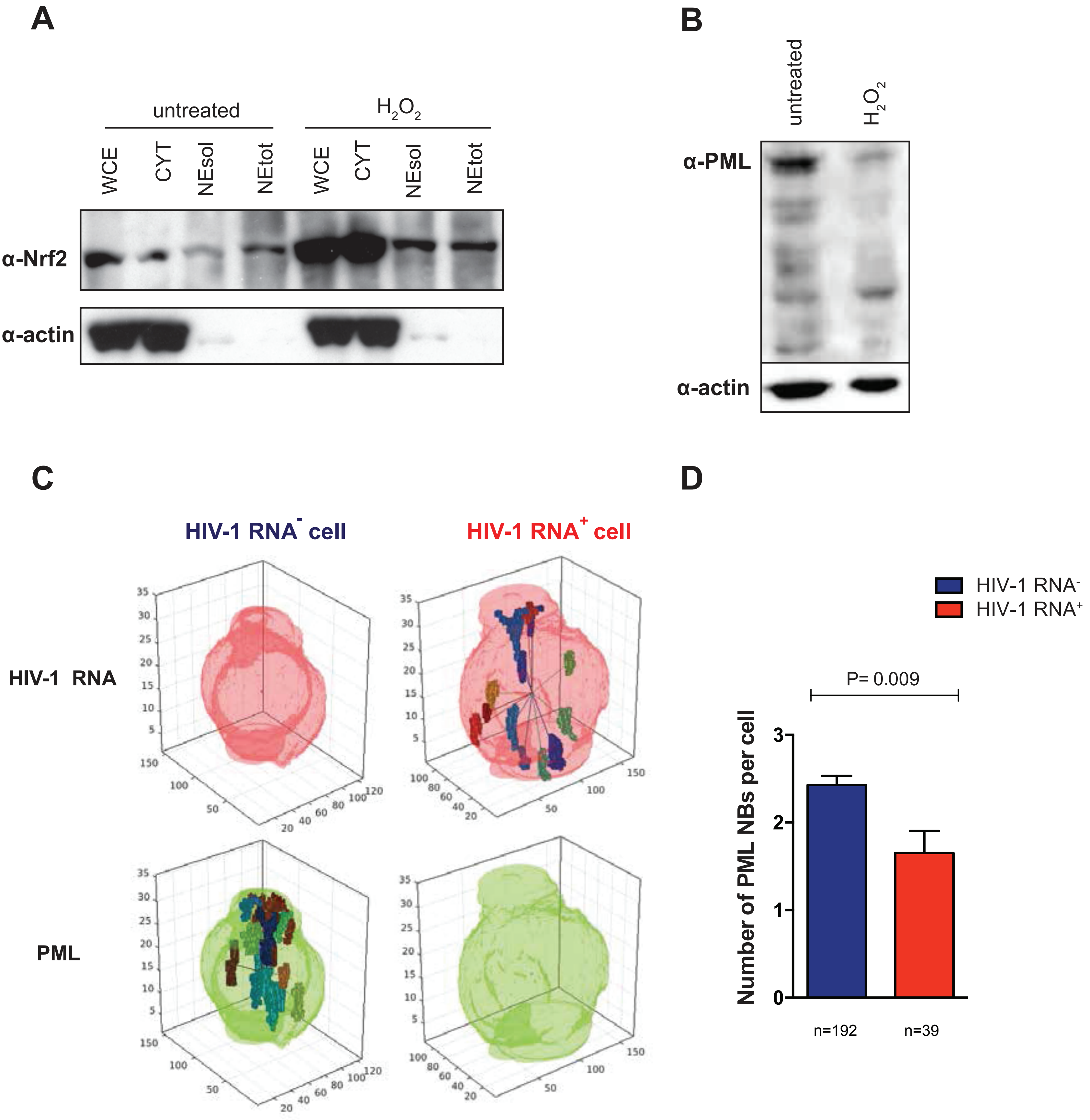

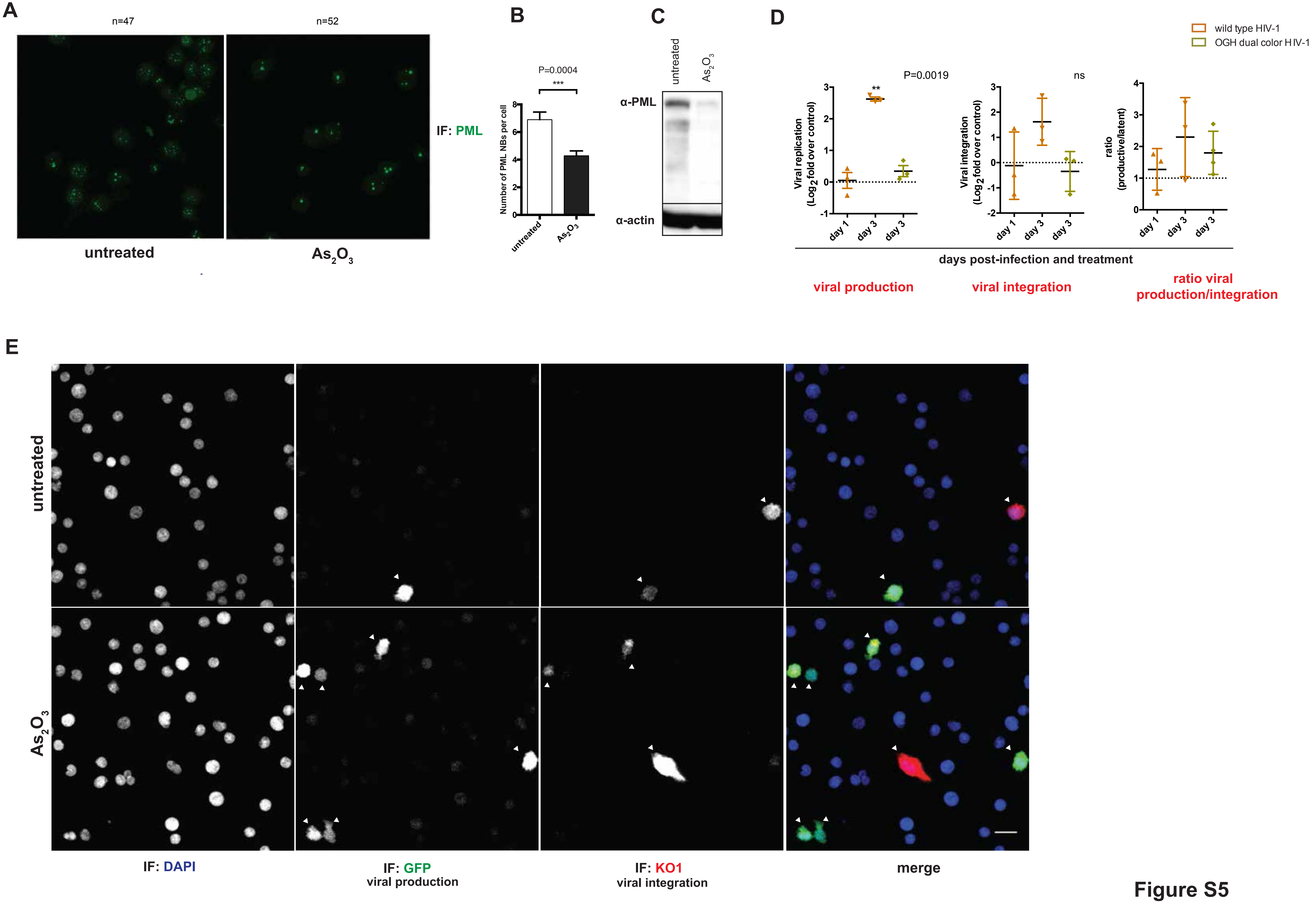

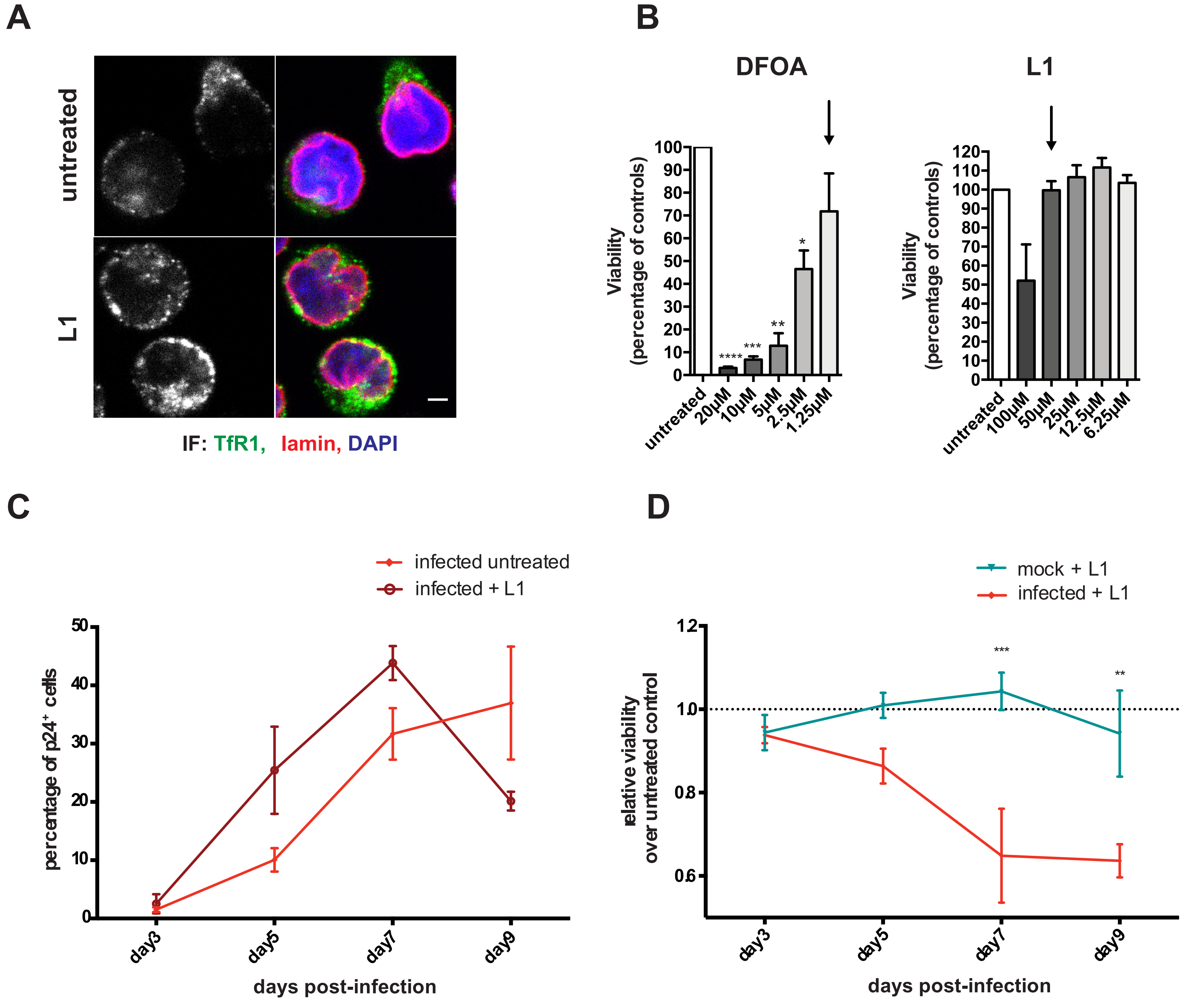
